# FAST RESENSITIZATION OF G PROTEIN-COUPLED RECEPTORS REQUIRES THEIR PI(4,5)P_2_-DEPENDENT SORTING INTO AN AP2 POSITIVE COMPARTMENT

**DOI:** 10.1101/2025.03.28.645988

**Authors:** Gergo Gulyas, Takashi Baba, Daniel J. Toth, Asuka Inoue, Tamas Balla

**Affiliations:** Section on Molecular Signal Transduction, Eunice Kennedy Shriver National Institute of Child Health and Human Development, National Institutes of Health, Bethesda, MD, USA; Department of Molecular Genetics, Wakayama Medical University, Wakayama, Japan; Department of Physiology, Semmelweis Medical University, Budapest, Hungary; Graduate School of Pharmaceutical Sciences, Tohoku University, 6-3, Aoba, Aramaki, Aoba-ku, Sendai, Miyagi, 980-8578 Japan

**Keywords:** GPCR desensitization, endocytosis, AT1 receptor, EFR3, PIP5K, arrestin, AP2, clathrin

## Abstract

G protein coupled receptors (GPCRs) are the target of about 35% of FDA-approved drugs, which underlines their importance in physiology and disease. Many GPCRs undergo rapid desensitization upon agonist activation, mediated by β-arrestin binding to the phosphorylated receptors, which prevents their G protein coupling. How such receptors regain their G protein signaling competence is poorly understood. Using the AT1 angiotensin II receptor (AT1R), we show that its rapid re-sensitization requires the sorting of the receptors into an AP2-positive plasma membrane (PM) compartment without receptors having to be internalized. This process requires the scaffolding protein, EFR3 and a dedicated PI(4,5)*P*_2_ pool specifically produced by PIP5KA. While β-arrestin 1 and −2 both can carry the receptors to the AP2 compartment, weaker binding of β-arrestin 1 to the receptor allows some of the receptors to re-sensitize, while strong β-arrestin 2 binding elicits stronger desensitization and directs the receptors for internalization. These results suggest that the rapid phase of re-sensitization of GPCRs occurs without their endocytosis and primarily takes place at the PM at specific steps during clathrin-coated pit maturation. Together with differences observed between the two β-arrestins to interact with CCPs and the receptors, our data suggest that specific PI(4,5)*P*_2_ pools controlled by EFR3A and PIP5K1A determine the balance between β-arrestin1 and −2 receptor interaction and delivery to the AP2 positive compartment, ultimately determining what fraction of the receptors regain their G protein signaling competence.

## INTRODUCTION

G protein coupled receptors (GPCRs) play fundamental roles in communicating environmental cues to the cell by generating intracellular signals. Most GPCRs respond to hormones or neurotransmitters with activation of heterotrimeric G-proteins that, in turn, trigger a cascade of downstream responses that are characteristic of the receptor, the G protein and the cell-type ^1^. Many activated receptors are subject to phosphorylation by G protein coupled receptor kinases (GRKs), or other protein kinases, such as PKA or PKC ^2,3^ followed by association with β-arrestins (β-arrestin 1 or −2) ^4^. β-arrestin binding to the receptor prevents its interaction with G proteins and, hence, uncouples the receptor from its primary signaling partner, a process termed desensitization ^5^. However, receptors bound to β-arrestins still can activate additional signaling pathways, thereby providing an added modality to their signaling properties ^6–9^. GPCRs that are bound to β-arrestins, are also subject to internalization primarily via clathrin-mediated endocytosis (CME), although parallel pathways of GPCR endocytosis have also been described ^10–13^. GPCR endocytosis effectively reduces the number of cell surface receptors and hence limits their ability to activate G proteins, although elegant recent data suggest that the internalized receptors still can signal from endocytic compartments ^14–16^. Such endocytosed receptors need to be recycled to the PM to regain their G protein signaling competence ^17–20^.

β-arrestins, therefore, are important both for the rapid uncoupling of the receptors from their partner G proteins and also for the removal of the receptors from the cell surface and directing them to the endocytic sorting machinery ^5^. β-arrestins also serve as adaptor molecules that sort receptors to clathrin-coated pits (CCPs) and interact with both the adaptor protein complex 2 (AP2) and clathrin, directing receptors to CCPs and ultimately sorting them to endocytic vesicles for further processing ^12,21^. During CME, receptors first converge by lateral diffusion within the plasma membrane into preformed, nascent CCPs that are assembled by multiprotein complexes, such as FCHO1 and −2, EPS15, EPS15L1 or AP180 ^22,23^. CCPs holding their cargoes further mature and gain an increasing membrane curvature and eventually bud from the PM by the sequential action of membrane deforming proteins, such as the endophilins and the fission promoting dynamins ^24–26^. The destination of the internalized vesicles holding the receptors will depend on their protein interacting partners and the properties of the GPCR-arrestin complex itself. In this respect, GPCRs are classified into two groups: Class A GPCRs, such as their prototype β-adrenergic receptors, show relatively weak affinity to β-arrestins resulting in the quick release of the arrestins from the receptors ^12,27^. In contrast, Class B receptors, such as the vasopressin type 2 (V2R) or the AT1R maintain tight β-arrestin binding and the receptor-β-arrestin complex travels through the endocytic machinery either directing the receptor for degradation in lysosomes or for recycling back the PM ^28–31^. It is generally believed, although not firmly established, that re-sensitization of GPCRs requires dissociation of the GPCR-β-arrestin complex following dephosphorylation of the receptors by phosphoprotein phosphatases, a process that occurs in the endocytic compartments ^18^. Whether β-arrestins can dissociate from the receptor in the PM and, if so, how this process is regulated, are questions that are relatively poorly understood.

Phosphatidylinositol 4,5-bisphosphate (PI(4,5)*P*_2_) and its precursor phosphatidylinositol 4-phosphate (PI4*P*) have been known as phospholipase C (PLC) substrates and the precursors of second messengers generated by GPCRs that activate the PLC enzymes ^32^. However, PI(4,5)*P*_2_ in the PM also interacts directly with several GPCRs ^33^ as well as other proteins that affect GPCRs, such as GRK2 ^34,35^ and, as shown most recently, β-arrestins ^22,36–38^. PI(4,5)*P*_2_ is also critical for the endocytosis process via direct interaction with proteins of the endocytic machinery, such as AP-2, Epsin, AP180 and dynamin ^39^.

Given this multitude of PI(4,5)*P*_2_ targets, the question arises, how all of these processes affect the properties and fate of GPCRs during their stimulation, particularly, in the case of receptors that activate PLC. In the present study we investigated the question of how desensitized GPCRs regain their G protein coupling competence. Specifically, we addressed the contribution of PI(4,5)*P*_2_ to the control of GPCR G protein coupling using the AT1 angiotensin II receptors (AT1R), which use PLCβ activation as their primary signaling output. Our studies reveal that a fraction of the AT1Rs can regain their G-protein coupling competence without internalization but require sorting to CCPs by a process that is controlled by a specific form of PIP 5-kinase, the scaffolding protein, EFR3, and is differentially affected by the two β-arrestins as they bind to the receptors.

## RESULTS

### PIP5K1A regulates G protein coupling of GPCRs

AT1Rs are coupled to PLCβ activation through the heterotrimeric G_q/11_ family of proteins resulting in the rapid hydrolysis of PI(4,5)*P*_2_ to Ins(1,4,5)*P*_3_ and diacylglycerol ^40,41^. During this process, PI(4,5)*P*_2_ pools are maintained by the sequential actions of the PI 4-kinase, PI4KA and the type I PIP 5-kinases (PIP5K1) (Fig. 1A). PI4KA works in the PM as part of a ternary molecular complex composed of PI4KA, FAM126A, TTC7 and EFR3, the latter providing PM anchoring ^42,43^. In mammalian cells, EFR3 proteins exist in two highly similar isoforms (A and B). In previous studies, we showed that silencing of EFR3 promotes receptor phosphorylation and facilitates receptor desensitization ^44^. This finding prompted us to further investigate whether PI(4,5)*P*_2_ and specifically, the PIP5K1 enzymes play any role in the control of receptor desensitization. This question was especially relevant in GPCRs that are coupled to PLC activation and cause PI(4,5)*P*_2_ hydrolysis.

**Figure 1.**
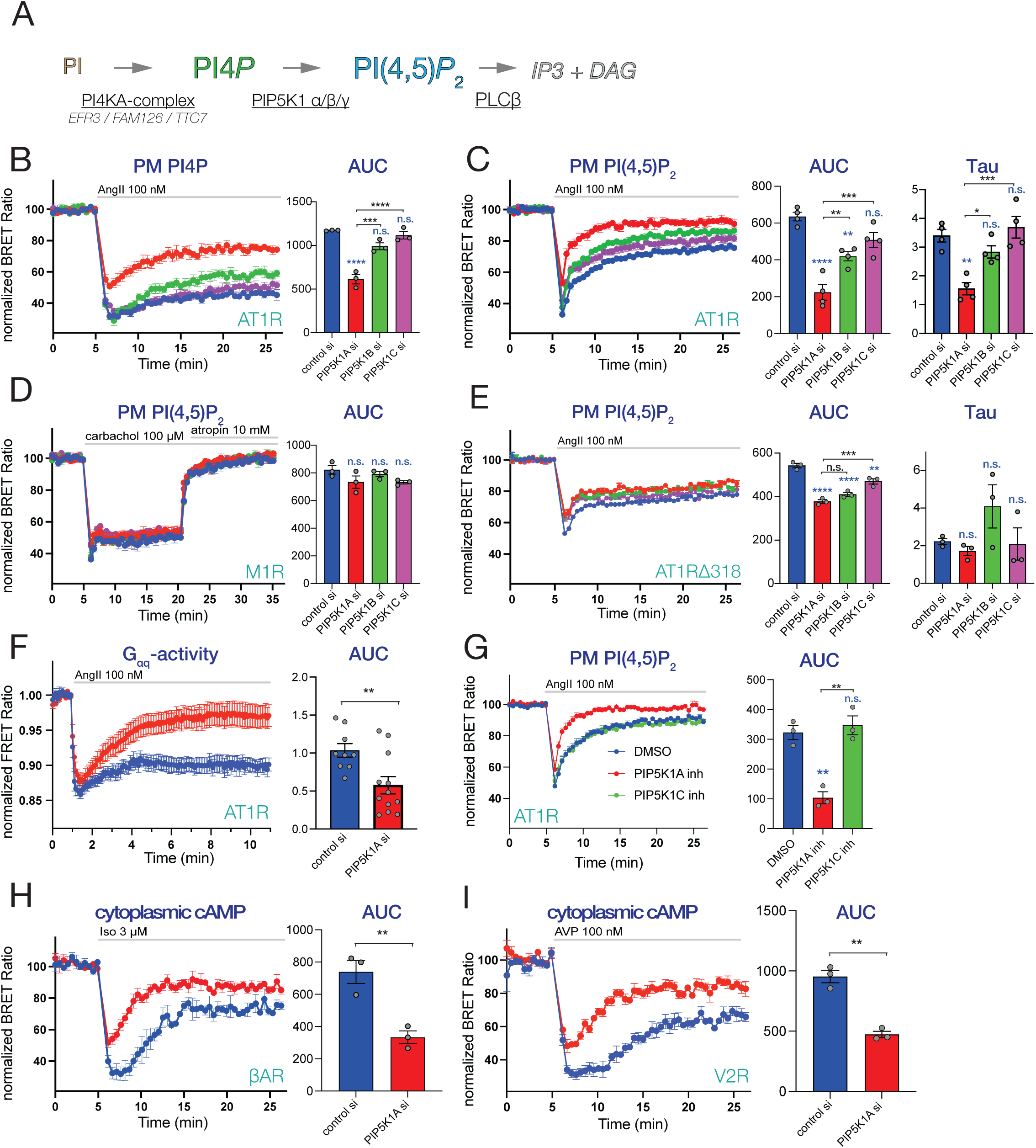
The contribution of phosphatidylinositol 4-phosphate 5-kinases (PIP5Ks) to the maintenance of GPCR activity. **(A)** A simplified cartoon showing that phosphatidylinositol 4,5-bisphosphate [PI(4,5)P_2_] is generated in subsequent enzymatic steps from phosphatidylinositol (PI), catalyzed by phosphatidylinositol 4-kinase alpha (PI4KA) and phosphatidylinositol 4-phosphate 5-kinases (PIP5K1α/β/γ). PI4KA works as a ternary complex formed with TTC7 and FAM126 proteins, anchored to the PM with the EFR3A or EFR3B proteins. Gq-coupled GPCRs activate PLCβ at the PM to generate the two second messengers, inositol 1,4,5-trisphosphate (IP_3_) and diacylglycerol (DAG) from PI(4,5)P_2_. **(B-E)** Agonist-induced changes in PM PI4*P* (B) and PI(4,5)P_2_ (C-E) levels monitored by BRET analysis in cells depleted in the various PIP5K1A isoforms (blue-control; red-PIP5K1A; green-PIP5K1B; purple-PIP5K1C). HEK-AT1R (B-D) and HEK-AT1R-Δ318 (E) cells were transiently transfected with the respective BRET-probe alone (B, C and E), or together with muscarinic type1 (M1R)(D). After a 5 min control period, cells were treated with 100 nM angiotensin II (AngII) (B-C and E) or 100 μM carbachol (CCh) followed by the addition of 10 μM atropine after 15 mins (D). Bar graphs next to the curves show area under the curve (AUC) calculated as deviations from the baseline during the stimulated period or association rate constants (Tau) calculated from the peak depletion point. Curves show the means ± SEM of three (B, D and E) or four (C) independent experiments, each performed in triplicates. The individual experimental data are shown as scatter plots in the AUC and Tau plots. One-way ANOVA followed by Tukey post-hoc test was used to estimate differences between the separate groups in multiple comparisons. Blue asterisks show comparisons to control, while black asterisk refer to the indicated comparisons. (n.s.: non-significant; *: p < 0.05; **: p < 0.01; ***: p < 0.001; ****: p < 0.0001). **(F)** Measurements of Gq-activity in control (blue) and PIP5K1A-depleted (red) HEK-AT1 cells using a FRET-based Gq biosensor. Cells transfected with the Gq biosensor for 24 h, were treated with 100 nM AngII and FRET ratio values, normalized to the initial resting values (I/I_0_) were plotted (see Methods for details). AUC calculations are shown next to the curves to compare the two groups. Statistical significance was obtained using two tailed, unpaired t-test. Scatter plots show the average of recorded cells in individual experiments. Data are means ± SEM obtained from nine (control) and twelve (PIP5KIA KD) dishes (5-8 cells per dish) in three independent experiments (**: p < 0.01) **(G)** BRET measurements showing PM PI(4,5)P_2_ changes in HEK-AT1R cell pretreated with DMSO (blue) or with the indicated inhibitors: 10 nM PMA-31 to inhibit *PIP5K1A* (red curve) or 500 nM UNC3230 to inhibit *PIP5K1C* (green curve) for 30 min before addition of 100 nM AngII. Bar graphs show AUC calculations. One-way ANOVA followed by Tukey post-hoc test for multiple comparison was used to assess statistical significance between the groups. Scatter plots show the results of individual experiments. The curves show means ± SEM of three independent experiments, each performed in triplicates. NS designates no statistical difference compared to controls, while black asterisks indicate individual comparisons. (**: p < 0.01). **(H-I)** Monitoring changes in cytoplasmic cAMP levels in HEK-AT1R (H) or HEK-V2R (I) cells stimulated with 3 μM isoprenaline to activate endogenous β-adrenergic receptors (βAR) or with 100 nM AVP to stimulate stably expressed human vasopressin type-2 receptors (V2R). Cells were treated with control siRNAs (blue) or siRNAs targeting PIP5K1A (red) and subsequently transfected with a BRET-based cAMP-sensor. Bar graphs show AUC calculations to compare the respective groups using two tailed, unpaired t-test. Scatter plots show the results of individual experiments. The curves show means ± SEM of three independent experiments, each performed in triplicates. (**: p < 0.01).

In mammalian cells, PIP5K1 enzymes exist in three forms, α, β and γ with five splice variants of PIP5K1C ^45,46^. To assess the role of individual PIP5Ks, we knocked-down (KD) the enzymes using isoform-specific siRNA pools in a HEK293 cell line that stably express the AT1R (HEK-AT1) ^47^. Western blot analysis showed that PIP5K1A and PIP5K1C were the dominant forms over the PIP5K1B as detected by the antibodies we used, and the efficiency of KD of the individual enzymes (Fig. S1A). We monitored PI4*P* (Fig. 1B) and PI(4,5)*P*_2_ levels (Fig. 1C) in the PM during AngII stimulation using our previously described lipid biosensors in BRET analysis ^48^. Importantly, none of the siRNA pools targeting individual PIP5Ks altered the basal levels of PI4*P* or PI(4,5)*P*_2_ based on basal (unstimulated) BRET ratios (Fig. S1B and C). This was not surprising given the redundancy between the PIP5K1 proteins in PI(4,5)*P*_2_ maintenance ^49^. Strikingly, KD of PIP5K1A, but not the other two PIP5Ks, had a robust effect on both PI4*P* and PI(4,5)*P*_2_ recoveries, during AngII stimulation, causing a faster recovery of both of these lipids compared to controls (Fig. 1B and C). This result was surprising, as silencing of PIP5K enzymes is expected to slow down rather than speed up the rate of PI(4,5)*P*_2_ resynthesis. This finding raised the possibility that KD of PIP5K1A enhanced the desensitization of the AT1R causing the faster recovery of the PM PI4*P* and PI(4,5)*P*_2_ levels during AT1R activation. To test this possibility, we carried out similar knockdown experiments in HEK293 cells that expressed either the M1 muscarinic acetylcholine receptors (M1R) or a phosphorylation deficient C-terminally truncated mutant AT1R (Δ319). The M1R receptor is known for very slow desensitization ^50,51^, and the truncated AT1R is unable to desensitize ^52^. The other advantage of using the M1 receptor is that its activation by agonist (carbachol) can be rapidly reversed by atropine, which allows assessment of the PI(4,5)*P*_2_ resynthesis process. These experiments showed that PIP5K1A KD did not affect PI(4,5)*P*_2_ (Fig. 1D) or PI4*P* (Fig. S1D) recovery when the M1 receptors were stimulated first by carbachol (100 μM) followed by reversal with atropine (10 μM). Similarly, no difference in PI(4,5)*P*_2_ kinetics was observed when truncated AT1Rs were used to stimulate PLCs (Fig. 1E). [Note that the rapid immediate rise in PI(4,5)P_2_ signal after its peak depletion is a reflection of the transient IP_3_ peak that contributes to the displacement of the PLCδ1PH from PI(4,5)P_2_ ^53,54^. While KD of the PIP5Ks had a small effect in reducing the size of PI(4,5)*P*_2_ drop after AngII stimulation, they did not affect the slow rate of recovery (Fig. 1E). These data taken together suggested that the faster recovery of these PIP lipids observed in PIP5K1A depleted cells during stimulation of the AT1Rs was indeed related to the faster desensitization of the AT1Rs.

Next, we used an alternative approach to directly monitor the G-protein coupling activity of the AT1R by following Gq dissociation caused by receptor stimulation employing a FRET-based Gq sensor ^55^. These experiments also confirmed the more transient Gq activation in cells where PIP5K1A was depleted (Fig. 1F). Additionally, we turned to a pharmacological approach to inhibit PIP5Ks using a recently described PIP5K1A inhibitor ^56^, (PMA-31, kindly provided by Ehab El-Awaad from the Joachim Jose laboratory) and the PIP5K1C-specific inhibitor, UNC3230 ^57^, and tested their effects on the PI(4,5)*P*_2_ response after stimulating wild type AT1Rs. These results showed that PIP5K1A but not PIP5K1C inhibition accelerated PI(4,5)*P*_2_ recovery during AngII stimulation (Fig. 1G), essentially confirming the findings of the knock-down experiments.

Next, we investigated whether PIP5K1A gene silencing affects the coupling efficiency of other GPCRs that do not primarily activate PLCs. β-adrenergic receptors (βARs) and vasopressin type-2 receptors (V2R) are known to activate the Gs signaling pathway leading to increased cytosolic cAMP levels. HEK293 cells endogenously express βARs, whereas for the investigation of V2R responses, we generated a V2R expressing stable cell line. Cytoplasmic cAMP changes were assessed using the previously described BRET-based cAMP sensor ^58^. These experiments showed that PIP5K1A KD also resulted in a faster cAMP recovery after isoprenaline or AVP stimulation (Fig. 1H and I, respectively).

Based on these results, we concluded that a PI(4,5)*P*_2_ pool, specifically made by PIP5K1A, has a unique contribution to support the rapid re-sensitization of the tested GPCRs regardless of their G protein coupling preference.

### EFR3A knock down phenocopies PIP5K1A silencing, and promotes accelerated GPCR-desensitization

In a previous work, we showed that combined silencing of EFR3A and -B caused AT1R hyperphosphorylation and a faster desensitization during agonist stimulation ^44^. To determine whether EFR3 isoforms differ with respect to their importance for receptor re-sensitization, we knocked down the two isoforms independently and followed PI(4,5)*P*_2_ and PI4*P* kinetics after AngII stimulation (Fig. 2A and Fig. S1E, respectively). We found that KD of EFR3A was more effective in promoting faster desensitization of AT1Rs than that of EFR3B. The effect of EFR3A KD on AT1R-mediated Gq-protein activation was also monitored using the FRET-based approach, yielding similar effects of EFR3 silencing on AT1R desensitization (Fig. 2B). Remarkably, the effect was opposite in the case of the βAR receptors, where EFR3B silencing resulted in a faster desensitization of βAR than silencing of EFR3A (Fig. 2C). V2R stimulation, on the other hand was affected by KD of either of the EFR3s (Fig. 2D).

**Figure 2.**
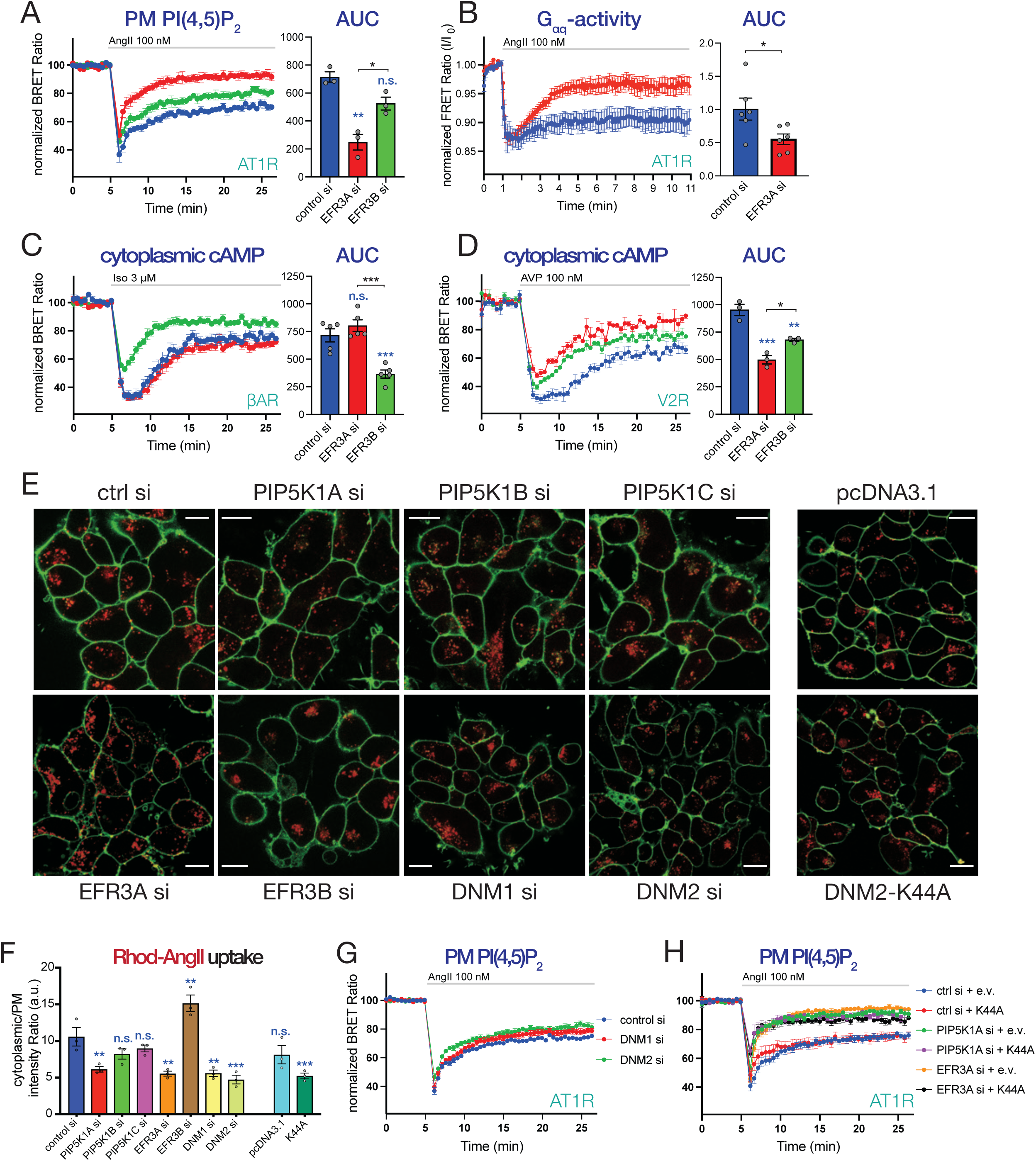
Effects of knockdowns of PIP5K1A or EFR3 on GPCR activity and internalization. (**A**) BRET analysis of PM PI(4,5)P_2_ changes in HEK-AT1 cells depleted in either EFR3A (red) or EFR3B (green) and stimulated with 100 nM AngII. HEK-AT1 cells were treated with the indicated siRNAs for 4 days (total) and transfected with the PI(4,5)P_2_ BRET sensor one day before BRET analysis. Bar graphs show AUC calculations to compare the treated groups using one-way ANOVA followed by Tukey post-hoc test for multiple comparison. Scatter plots show the results of individual experiments. Curves are shown as means ± SEM of three independent experiments, each performed in triplicates. Blue asterisks show comparisons to control, while black asterisk refer to the indicated comparison. n.s. indicates no significant difference compared to control. (*: p < 0.05; **: p < 0.01). (**B**) Monitoring Gq-activity in control (blue) and EFR3A (red)-depleted HEK-AT1 cells using a FRET-based Gq-biosensor. Following EFR3A knockdown, cells were transfected with the Gq-biosensor for 1 day and stimulated with 100 nM AngII. Fluorescence was monitored in both the YFP and CFP channels using excitation at 410 nm. FRET Ratio values were calculated simply forming a YFP/CFP ratio for each cell followed by normalization to the initial resting ration values (I/I_0_). Bar graphs show AUC calculations for statistical comparison of the two groups using two tailed, unpaired t-test. Scatter plots show the calculated AUC values from each of six individual dishes (8-10 cells per dish) recorded in 3 independent experiments (*: p < 0.01). (**C**) βAR and **(D)** V2R activity were monitored as changes in cytoplasmic cAMP levels using a BRET-based cAMP sensor. Recordings form control (blue), EFR3A (red)- or EFR3B (green) -depleted HEK-AT1 (C) or HEK-V2R cells (D) are shown. Cells were stimulated with 3μM isoprenaline (Iso) (C) or 100 nM vasopressin (AVP) (D). Bar graphs show AUC values calculated in individual experiments and analyzed by one-way ANOVA to compare the groups and to calculate statistical differences. Graphs show means ± SEM of five (C) and three (D) independent experiments, each performed in triplicates. Blue asterisks represent comparisons to controls, while black asterisk show comparisons between the indicated groups (NS.: non-significant; *: p < 0.05; **: p < 0.01; ***: p < 0.001). *Please note that the control curve presented in panel (D) is identical to the one displayed on* Fig. 1I. *Both the PIP5K1A and EFR3 knock-down experiments were executed at the same time and on the same plate in parallel recordings*. (**E**) Representative confocal images showing internalization of Rhod-AngII (red) in HEK-AT1R cells treated with the indicated siRNAs or transfected with pcDNA3.1 (as control) or dominant negative Dynamin2 (DNM2-K44A). Green fluorescence highlights the plasma membrane stained with CellMask. Scale bar represents 10 μm. (**F**) Quantification of image sequences as shown in panel (E). Bar graphs show intensity ratios of Rhodamine fluorescent intensities between the cytoplasm and the PM. Data are means ± SEM obtained in three independent experiments, and analyzing at least three captured fields of view for each treatment in each experiment. Scatter plot shows individual ratio values calculated in each separate experiments. Statistical significance was evaluated using ANOVA, followed by Tukey post-hoc test. Blue asterisks refer to comparisons to the control. (n.s.: non-significant; **: p < 0.01, ***: p < 0.001). (**G**) Monitoring PM PI(4,5)P_2_ levels using BRET analysis in HEK-AT1R cells after knocking down Dynamin1: (DYN1-red) or Dynamin2: (DYN2 - green) upon 100 nM AngII stimulation. Data are means ± SEM of five independent experiments, each performed in triplicates. (**H**) Monitoring PM PI(4,5)P_2_ levels in AngII-stimulated HEK-AT1R cells that express either an empty vector (e.v.) or a dominant negative form of Dynamin2 (K44A) following treatment with the indicated siRNAs.

Notably, both PI(4,5)*P*_2_ recovery after AngII stimulation and cAMP kinetics after isoproterenol treatment were similarly affected by knock-down of either PIP5K1A or the EFR3s (Fig. S1F and G, respectively). Double KD experiments showed no additivity between the effects of EFR3A and PIP5KA, suggesting that these two proteins may act on the same pathway that controls the desensitization of the AT1R and βARs (Fig. S1F and G).

#### Internalization is not required for the rapid phase of GPCR re-sensitization

PM PPIns are important regulators of the GPCR internalization process, and massive reduction of PM PI(4,5)*P*_2_ reduces GPCR internalization ^59^. It has been suggested that internalization of GPCRs and their subsequent recycling to the PM serves as a re-sensitization process ^60^. According to these models, the effect of the KDs of either PIP5K1A or EFR3A, could exert their effects by preventing the internalization of the receptors. To test the effects of EFR3 or PIP5KA KDs on AT1R internalization, we followed the uptake of fluorescently labeled AngII (Rhod-AngII). Using confocal images (Fig. 2E), we quantified the fluorescent signal at the PM (using a PM mask) and inside the cells and calculated a cyto/PM intensity ratio (Fig. 2F). These experiments showed, that indeed, the KDs of either EFR3A or PIP5KA resulted in a reduced AngII intracellular uptake (represented by lower cytoplasmic/PM fluorescent intensity ratios) compared to those measured in control siRNA treated cells (Fig. 2F) (note that not every cell is expected to be efficiently “knocked down” in these fields of view). However, depletion of Dynamin 1 and 2 or overexpression of the dominant negative Dynamin2 (DNM2-K44A) ^61,62^, while comparably decreased the uptake of AngII (Fig. 2E and F), they failed to affect the kinetics of PI(4,5)*P*_2_ after AngII stimulation (Fig. 2G-H and Fig. S1H-J). Although silencing of Dynamin2 resulted in a slight reduction in the drop of PM PI(4,5)*P*_2_ levels reflected in the AUC calculations (Fig. S2H), it did not affect the rate of PI(4,5)P_2_ recoveries as indicated by the Tau calculations (Fig. S2I). These data collectively suggested, that while both PIP5KA and EFR3 affect molecular events important for receptor internalization, their effects on AT1R re-sensitization are upstream of dynamin function and may not involve internalization of the receptors. It is worth pointing out, here, that GPCR desensitization also results from the reduction of the number of receptors from the cell surface, also termed down-regulation ^5,63,64^. This latter process is tightly coupled to receptor internalization and for those internalized receptors to regain their G-protein coupling competence, receptors must be recycled. Our data do not argue against the importance of this process that occurs on a slower timescale.

#### PIP5K1A and EFR3A knock-down increases ß-arrestin2 and reduces ß-arrestin1 PM recruitment

Based on our data, the faster desensitization of AT1Rs observed after PIP5K1A or EFR3A silencing required the tail of the receptor that is phosphorylated and binds β-arrestins. Therefore, we turned to the question whether β-arrestin function is altered by PIP5K1A or EFR3A KDs. This question was even more relevant in view of recent studies that showed the importance of PPIns in binding β-arrestins to GPCRs ^37,38,65^). First the PM recruitment of β-arrestin1 or −2 was tested using TIRF microscopy after knocking-down PIP5K1A or EFR3A. Our initial experiments revealed that the levels of β-arrestin expression from a CMV promoter-driven construct caused massive desensitization of the AT1Rs. Therefore, we utilized thymidine-kinase (TK) promoter-driven mRFP-tagged β-arrestins that allowed monitoring β-arrestin movements at extremely low expression levels. Silencing either PIP5K1A or EFR3A reduced PM recruitment of β-arrestin1 and made it more transient after AngII stimulation (Fig. 3A, 3D and Fig. S2A), whereas they increased β-arrestin2 recruitment (Fig. 3B, 3E and Fig. S2B). In parallel experiments, GFP-tagged AT1R clustering in response to AngII in PIP5K1A or EFR3A KDs showed an increase and was more sustained compared to the RNAi controls, consistent with the decreased receptor internalization process (Fig. 3C, 3F, and Fig. S2C). As the two β-arrestin isoforms compete for GPCR binding, our data suggested that PIP5K1A or EFR3A silencing shifts the balance toward β-arrestin2 binding over β-arrestin1.

**Figure 3.**
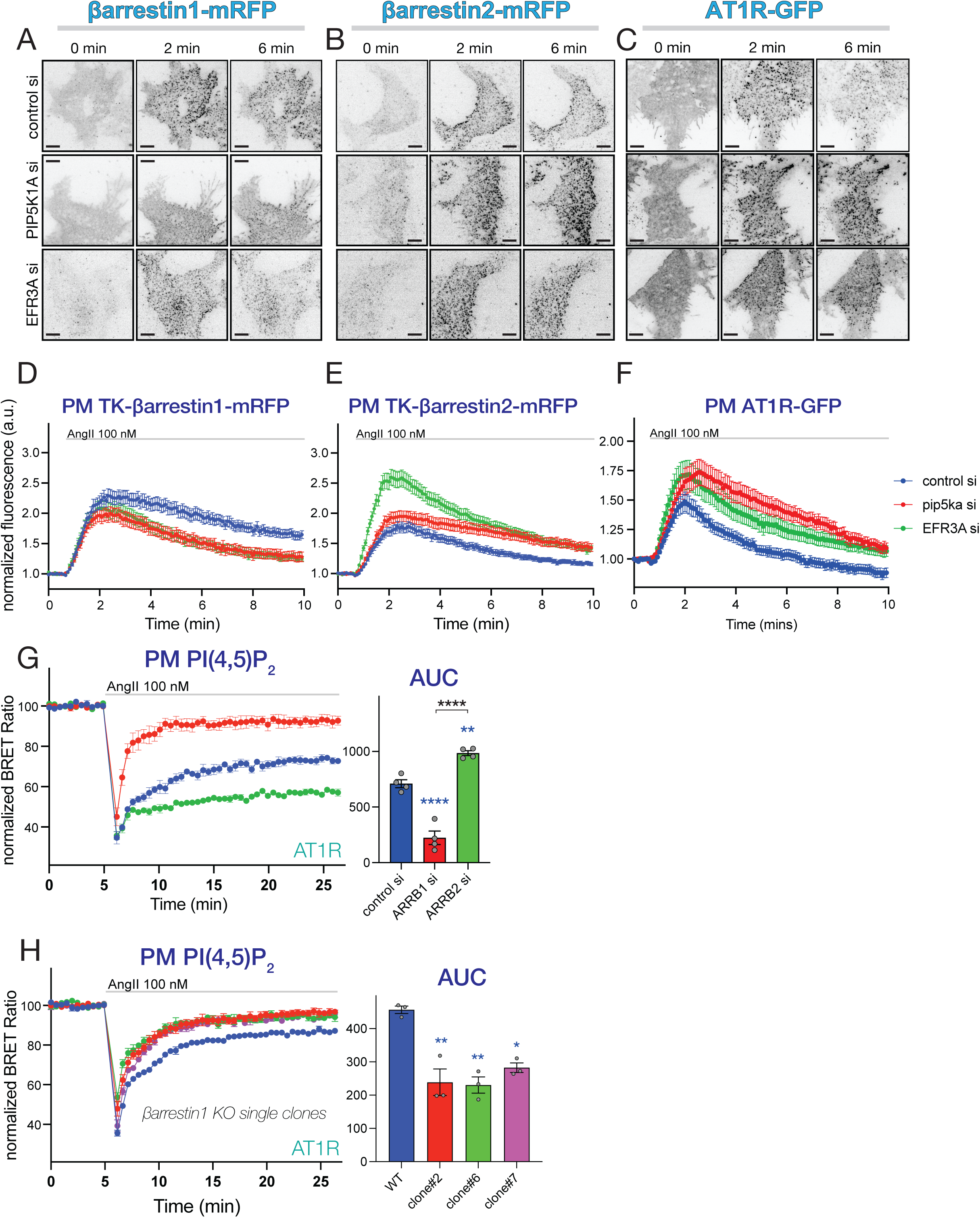
TIRF analysis of clustering the individual β-arrestins and the AT1R upon stimulation with AngII. **(A-C)** HEK-AT1cells were transfected with TK-promoter driven mRFP-tagged rat β-arrestin1 or β-arrestin2, after treatment with the indicated siRNAs to deplete PIP5K1A or EFR3A proteins. Cells were stimulated with AngII and clustering of the β-arrestins was monitored in a TIRF microscope (0, 2 and 5 min pictures are shown from a representative time-lapse recording). AT1R clustering was also monitored in HEK293 cells stably expressing the C-terminally GFP-tagged AT1R (C). Scale bar represents 10 μm. **(D-F)** Quantification of fluorescence intensities from several recordings similar to what is shown in panels A-C. Normalized fluorescence were calculated from average cellular intensities normalized in each case to the average intensity based on the last 3 images before stimulation. Note that the fluorescence intensities increase as clustering develops. Curves represent means ± SEM of data collected from 38 (control siRNA), 37 (PIP5K1A siRNA) and 31 (EFR3A siRNA) cells each in 13 independent dishes in 7 experiments (**D**), 51 (control siRNA), 43 (PIP5K1A siRNA) and 54 (EFR3A siRNA) cells in 14 (control siRNA and PIP5K1A siRNA) or 15 (EFR3A siRNA) independent dishes in 7 experiments (**E**) or 52 (control siRNA), 39 (PIP5K1A siRNA) and 46 (EFR3A siRNA) cells in 12 (control siRNA) or 9 (PIP5K1A siRNA and EFR3A siRNA) independent dishes in 5 experiments (**F**). **(G)** Monitoring PM PI(4,5)P_2_ levels in AngII-stimulated HEK-AT1 cells treated with control (blue), β-arrestin1 (red) or β-arrestin 2 (green) specific siRNA pools. Curves show means ± SEM of four independent BRET experiments, each performed in triplicates. Bar graphs show AUC values calculated for each of the experiments in the scatter plot. Multiple comparisons between the different groups were assessed by one-way ANOVA followed by Tukey post-hoc test. Blue asterisks refer to comparisons to control, while black asterisks refer to comparisons between the indicated groups. (**: p < 0.01; ****: p < 0.0001). **(H)** Changes in PM PI(4,5)P_2_ levels after AngII stimulation in selected β-arrestin1 knockout single clones using BRET analysis. Curves shown are means ± SEM of three independent experiments, each performed in triplicates. Bar graphs show AUC values calculated from each individual experiments and presented as scatter plots. One-way ANOVA followed by Dunnett’s multiple comparison was used to assess statistical differences. Significance levels refer to comparisons to WT cells (n.s.: non-significant; *: p < 0.05; **: p < 0.01).

### β-arrestin1 alleviates, whereas β-arrestin2 promotes AT1R desensitization and clustering

Next, we investigated how β-arrestin silencing affects AT1R G protein-coupling activity. As expected, β-arrestin2 KD resulted in the complete elimination of the desensitization of the AT1R (Fig. 3G, green). In striking contrast, silencing of β-arrestin1 had the opposite effect, namely it increased rather than decreased desensitization (Fig. 3G, red). The kinetic of the faster PI(4,5)*P*_2_ recovery was similar to those observed after PIP5K1A or EFR3A silencing. This unexpected finding with β-arrestin-1 silencing prompted us to follow up in several further experiments. First, we used RNAi resistant forms of the β-arrestins driven by TK-promoter to achieve low expression levels, mimicking endogenous levels in rescue experiments. Using this approach, we observed a partial rescue by overexpressing the siRNA-resistant βarrestin1 (Fig. S2D).

To rule out potential off target effects of the siRNAs, we tested multiple siRNAs targeting β-arrestin1 either individually or in combinations (Fig. S2E and F). Several of these additional β-arrestin1 siRNAs also increased AT1R receptor desensitization. However, as revealed by Western Blot analysis, some of them also reduced the level of β-arrestin2 and, therefore, had a mixed effect on AT1R desensitization (Fig. S2E and F). Thirdly, we generated β-arrestin1-KO cell lines using Cas9/CRISPR mediated genome editing. After single clone selection and confirming by Western Blotting (Fig. 3I), we identified several β-arrestin1 knockout clones that displayed similarly accelerated AT1R desensitization than what was seen after β-arrestin1 KD (Fig. 3H). However, again, some of the isolated clones did not show accelerated AT1R desensitization, but again, their β-arrestin2 levels were also found reduced (Fig. S2G, H and I).

Taken together, these data suggest, that β-arrestin1 binding protects AT1Rs from desensitization that is primarily driven by β-arrestin2 binding. This conclusion was also in agreement with the finding that PIP5K1A or EFR3A depletion shifts the balance toward β-arrestin2 binding to the receptor on the expense of β-arrestin1 (shown in Fig. 3A-F).

#### siRNA screen reveals that AP2 is critical for proper AT1R re-sensitization

So far, our data showed that the faster desensitization of AT1R observed in cells silenced in PIP5K1A or EFR3A, while associated with impaired receptor internalization, acted at step(s) that preceded dynamin-mediated fission. In an earlier study, Antonescu et al. described the role PIP5K1A in CCP initiation and maturation ^39^. To investigate whether components of CCPs affect the AT1R desensitization process, we performed an siRNA mini-screen targeting proteins that play roles in the early, middle, and late-phase of CCP formation (Fig. 4A-D). We also targeted endophilins, that were found to mediate clathrin-independent fast endocytosis ^10^. We found that silencing of the proteins contributing to the early (EPS15, EPS15L1, FCHO1 and FCHO2) or late phase (Endophilin1-3, a.k.a. SHGL2, SHGL1 and SHGL3, similarly to Dyn1 and Dyn2 (see Fig. 2G)) of the CME maturation failed to affect AT1R desensitization (Fig. 4B and D, and Fig. S3A and C respectively). In contrast, knocking-down the μ-subunit of AP2 complex (AP2M1) resulted in a significantly faster desensitization kinetic (Fig. 4C) while KD of clathrin heavy-chain (CLTC) had a minimal effect compared to controls (Fig. 4C and see and Fig. S3B for statistics). Notably, the faster kinetic by AP2M1 knockdown was highly similar to those observed after silencing PIP5K1A, EFR3A or β-arrestin1. These data suggested that these four proteins might be functionally connected and help AT1Rs maintain their G protein coupling competence. It is important to point out that β-arrestin2 knockdown, prevented the faster desensitization of the AT1R even when any of those other proteins (PIP5K1A, EFR3A or β-arrestin1 or AP2) were depleted, indicating that the β-arrestin2 binding is the ultimate cause of the accelerated receptor desensitization (Fig. S3D-G).

**Figure 4.**
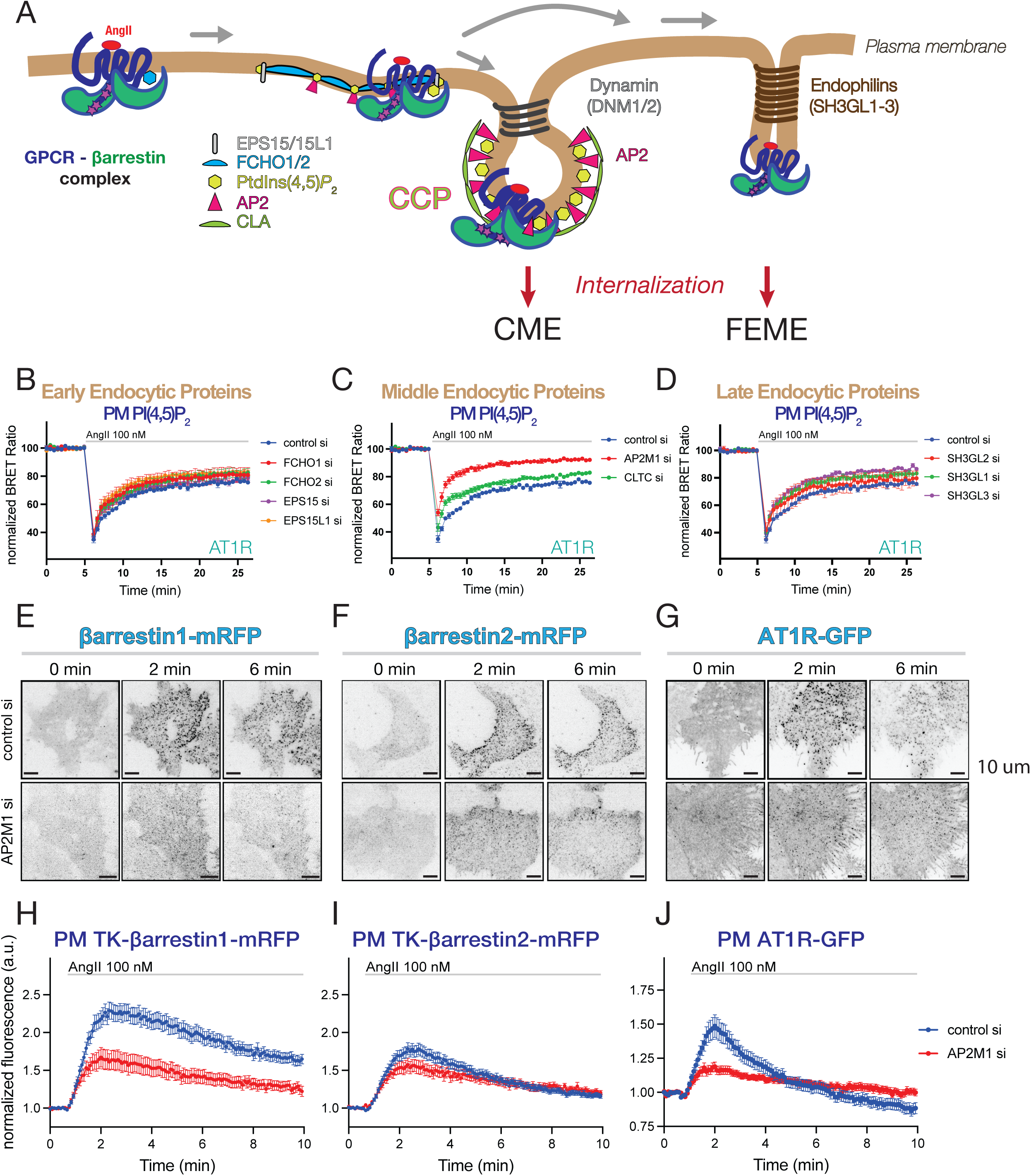
The AP2 adaptor of clathrin-coated pits is critical for fast AT1R receptor resensitization. **(A)** Simplified cartoon to illustrate important molecular players during maturation of clathrin coated pits (CCPs) leading to clathrin mediated endocytosis (CME). Endophilins have been identified as components of clathrin-independent, fast endophilin-mediated endocytosis (FEME) ^10^. **(B-D)** Plasma membrane PI(4,5)P_2_ changes assessed by BRET analysis in AngII-stimulated HEK-AT1 cells, pretreated with the indicated siRNAs to reduce the levels of molecules involved in early (B), middle (C) or late (D) phase of CME or FEME. Curves shown are means ± SEM of five (B-C) independent experiments. For curves shown in panel D, the number of experiments is four (SH3GL1 and SH3GL3 siRNAs) and three (SH3GL2 siRNA), each performed in triplicates. For statistics on AUC calculations, see Fig. S3 A-C. **(E-F)** Selected images from TIRF microscopy time-laps recordings show changes in β-arrestin1 and 2 PM recruitment and clustering after AngII treatment (0, 2 or 6 min) in HEK-AT1 cells transfected with TK-promoter driven mRFP-tagged rat β-arrestin proteins. One group of cells was treated with control siRNA or siRNA targeting the AP2 μ-subunit (AP2M1). **(G)** AT1R PM clustering and internalization following AngII stimulation (0, 2 or 6 min) monitored in HEK-AT1R-GFP cells by TIRF analysis and the effect of depletion of AP2 μ-subunits. *Please, note that for comparison, the same control cell pictures shown in Fig. 3A-C are also included in E-G.* Scale bars: 10 μm. **(H)** Quantification of recordings represented in ***(E)***. Values are average fluorescence intensities of cells, which were normalized to the average of the last 3 time points before stimulation. Curves represent means ± SEM of data collected from 38 (control siRNA – blue) and 25 (AP2M1 siRNA – red) cells in seven (control siRNA) or six (AP2M1 siRNA) independent experiments. **(I)** Quantification of recordings represented in ***(F)***. Values are average fluorescence intensities of cells, which were normalized to the average of the last 3 points before stimulation. Curves represent means ± SEM of data collected from 51 (control si – blue) and 29 (AP2M1 siRNA – red) cells in seven (control siRNA) or six (AP2M1 siRNA) independent experiments. **(J)** Quantification of recordings represented in ***(G)***. Values are average fluorescence intensities of cells, which were normalized to the average of the last 3 points before stimulation. Curves represent means ± SEM of data collected from 52 (control siRNA– blue) and 35 (AP2M1 siRNA – red) cells in five and three independent experiments. *Please note, that for comparison, the same control siRNA treated curves shown in* Figure 3 D-F *were also included in Figs H-I. (For further methodological details see Materials and Methods)*.

Next, we investigated how AP2 KD affected β-arrestin1 or −2 recruitment to the PM in TIRF experiments. For these experiments, again, we used TK-promoter-driven β-arrestin constructs to keep their expression level low (detected by the camera but often invisible by eye). We found that AP2 silencing significantly reduced PM clustering of β-arrestin1 (Fig. 4E and H) and both the clustering as well as the rate of internalization of AT1Rs (Fig. 4G and J). In contrast, AP2 KD had only a minor effect on β-arrestin2 PM clustering after AngII stimulation (Fig. 4F and I). These data suggested that AP2-depletion affects the ability of β-arrestin1, but not β-arrestin2 to be recruited to the PM clusters, and that both β-arrestin1 and −2 can deliver AT1Rs into the CCPs. Taken together, these experiments pointed to AP2 as an important hub in regulating β-arrestin1 function and controlling AT1R responsiveness in addition to its well-known role in CCP initiation, maturation and cargo sorting.

#### β-arrestin1 and β-arrestin2 show differences in their association with CCPs

The simplest conclusion from the data presented so far was that maintenance of AT1R G-protein coupling competence is related to the ability of β-arrestins (primarily β-arrestin1) to deliver the receptor to CCPs, by a process that requires AP2 and which is dependent on a PI(4,5)*P*_2_ pool, specifically generated by PIP5K1A with the help of EFR3 proteins. Moreover, there appeared to be a difference whether the receptors are sorted to the CCPs by β-arrestin1 or β-arrestin2. Given the recently recognized PI(4,5)*P*_2_ binding of the β-arrestins ^37,38,65^ and their different AP2 and clathrin-binding properties ^66–68^, it was important to analyze how the two β-arrestins interact with CCPs. For this, we turned to β-arrestin1/2 double knockout HEK293 cells ^69^, and simultaneously monitored clathrin light-chain (TK-mCherry-CLA), AT1R (AT1R-mVenus) and the two β-arrestin isoforms (TK-β-arrestin1/2-iRFP) in TIRF experiments. Using the co-localization analysis developed by the Taraska group ^70^, we found that β-arrestin2 showed co-localization with CCPs even before AngII stimulation, suggesting that β-arrestin2 is already prepositioned into CCPs even without binding to the receptor, since receptor clusters were not present in these CCPs without stimulation (Fig. 5A-D). Moreover, upon stimulation, β-arrestin2 showed a larger increase and co-localization with clathrin and brought the receptors more efficiently into the CCPs, than what was observed with β-arrestin1 (Fig. 5A-D).

**Figure 5.**
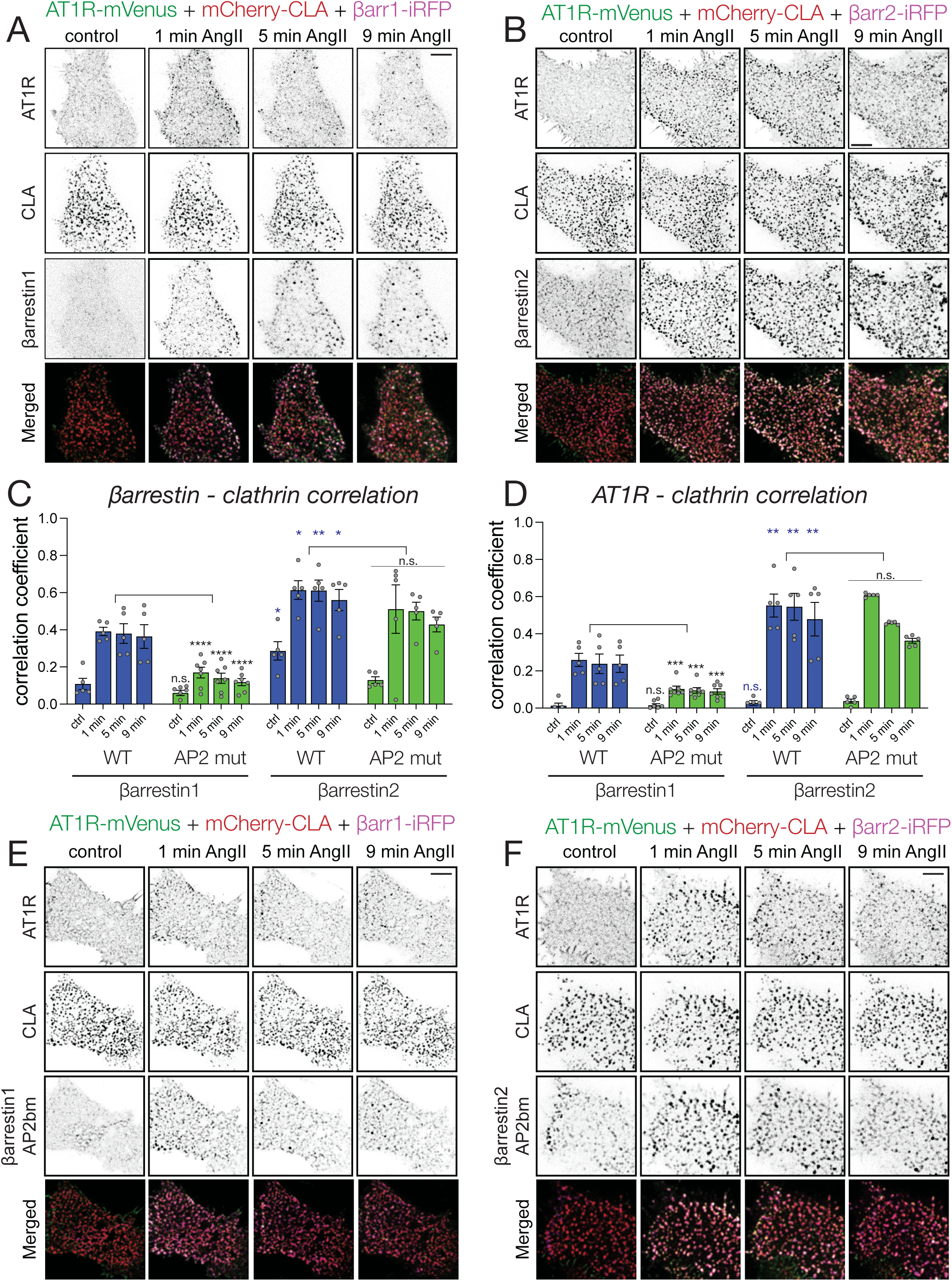
AP2-binding is important for βarr1 but not βarr2-mediated sequestration of AT1Rs into CCPs. **(A-B)** Representative TIRF microscopy images showing co-localization between angiotensin II receptor (AT1R-mVenus – green channel), clathrin (mCherry-CLA – red channel) and β-arrestin1 (TK-β-arrestin1-iRFP – magenta channel) **(A)** or β-arrestin2 (TK-β-arrestin2-iRFP – magenta channel) **(B)** in βarrestin1/2 double KO HEK293 cells after stimulating with 100 nM AngII. Scale bars: 5 μm. **(C-D)** Bar diagrams representing correlation coefficients calculated between clathrin and the indicated β-arrestins **(C)** or between clathrin and AT1R **(D)**. Blue and green bars correspond to wild type (WT) or AP2-binding mutant β-arrestin1 and −2 isoforms, respectively. Data are mean ± SEM of data collected from five (WT β-arrestin1, WT β-arrestin2 and AP2 mutant β-arrestin2) and seven (AP2 mutant β-arrestin1) cells based on analysis of hundreds of clathrin-coated structures on segmented images in each group. To compare the correlation values between β-arrestins1 and 2 (blue asterisks), or among wild-type (WT) and AP2 binding mutant (AP2 mut) arrestins in each group (black asterisks), two-way ANOVA followed by Holm-Šidak’s post hoc test was used. (n.s.: non-significant; *: p < 0.05; **: p < 0.01; ***: p < 0.001; ****: p < 0.0001) *(For further details see Materials and Methods)*. **(E-F)** Representative TIRF microscopy images demonstrating co-localization between angiotensin II receptor (AT1R-mVenus – green channel), clathrin (mCherry-CLA – red channel) and AP2-binding mutants of β-arrestin1 (TK-β-arrestin1-AP2bm-iRFP – magenta channel) **(E)** or β-arrestin2 (TK-β-arrestin2-AP2bm-iRFP – magenta channel) **(F)** in β-arrestin1/2 double KO HEK293 cells after stimulating with 100 nM AngII. Scale bars: 5 μm.

#### AP2 binding of β-arrestin1 but not of β-arrestin2 is necessary to direct and sequester AT1Rs into CCPs

After documenting the differences between the two β-arrestin isoforms, we decided to analyze further, how AP2 binding affects β-arrestins to recruit AT1Rs into CCPs. AP2-binding of β-arrestins occurs via their [DE]nX1–2FXX[FL]XXXR motif within their C-termini ^71,72^. In previous KD experiments we found that AP2-KD significantly affected the PM recruitment of β-arrestin1 but not of β-arrestin2 (Fig. 4C-F). To address this question in an alternative way, we performed similar co-localization analysis in cells that expressed mutant forms of β-arrestins consisting mutations in their related motif (R393A/R395A and R394A/R396A for β-arrestin1 and β-arrestin2, respectively) ^73^. In agreement with the conclusions of the AP2 KD experiments, we found that AP2 binding is necessary for β-arrestin1 (Fig. 5 C-E) to recruit and corral AT1R into CCPs, while it is less important in the case of β-arrestin2 (Fig. 5D-F).

#### AP2 binding of β-arrestin1 but not of β-arrestin2 limits its ability to desensitize AT1Rs

To analyze features of β-arrestins mutants with impaired AP2 interaction, we used the β-arrestin1/2 double knock out HEK293 cell line and overexpressed the β-arrestins together with the AT1R and the FRET-based Gq sensor and followed Gq activation using fluorescent lifetime imaging microscopy. Dissociation of the G protein βγ subunits tagged with Venus from the mTq-tagged α-subunit decreases the energy transfer between mTq and Venus ^55^ causing an increased lifetime of the mTq fluorescence. This analysis showed that wild-type β-arrestin2 (Fig. 6B and C) was more potent in desensitizing AT1Rs than the wild-type form of β-arrestin1. In the absence of β-arrestin2, overexpression of β-arrestin1 was also able to desensitize AT1Rs even if to a lesser extent (Fig. 6A and C). Introducing the AP2 binding mutations however, had a striking effect on β-arrestin1 (Fig. 6A and C) but not on β-arrestin2 (Fig. 6B and C) in that β-arrestin1-AP2-binding mutant was more effective in desensitizing the AT1R (Fig. 6A and C), similarly to wild type β-arrestin2 (Fig. 6B and C).

**Figure 6.**
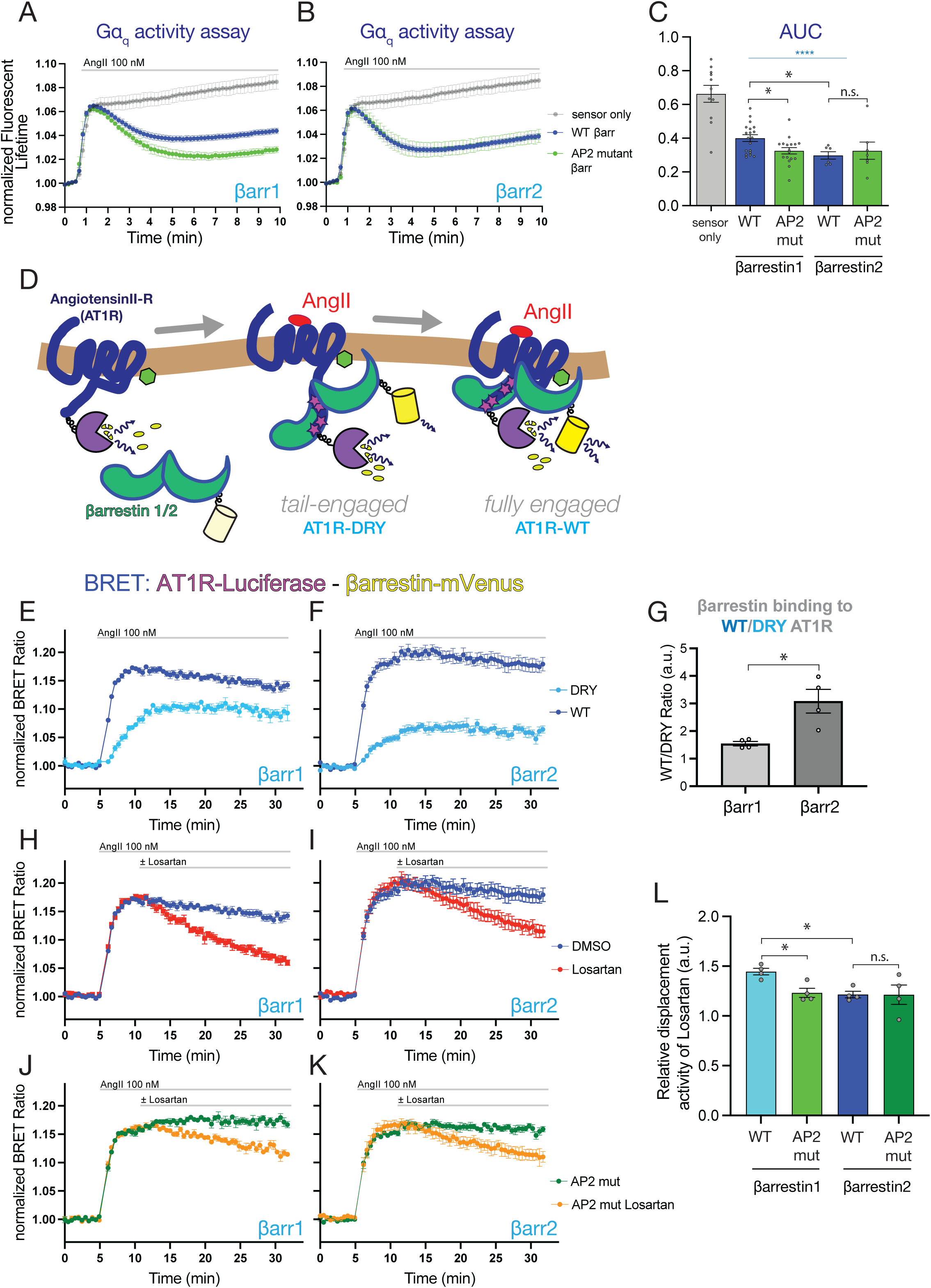
AP2-binding mutant β-arrestin1 but not β-arrestin2 is more potent in desensitizing AT1Rs than its wild-type form. **(A-B)** FLIM measurements show the effect of wild-type (WT) and AP2-binding mutants of β-arrestin1 **(A)** and β-arrestin2 **(B)** on G-protein activation by AT1Rs. Receptor activity was monitored with the FRET-based Gq sensor already described in Fig. 1A and B ^55^ except, that in these experiments, fluorescent lifetime of mTurqoise2 was used as a measure of FRET-efficiency. β-arrestin1/2 double KO HEK293 cells were co-transfected with untagged AT1R, the indicated mRFP-tagged β-arrestins and the Gq-sensor. Data were normalized to the 1″ values of control period before stimulation with 100 nM AngII. Data are means ± SEM of eleven (Gq), nineteen (β-arrestin1), seventeen (β-arrestin2), six (β-arrestin1-AP2 mut) and seven (β-arrestin2-AP2 mut) independent experiments. **(C)** Bar diagrams show the area under the curve (AUC) values calculated from the curves presented in (A and B). Data are means ± SEM from the number of experiments described for panels A and B. Statistical differences between the control cells (w/o β-arrestins) and those also expressing the β-arrestins were calculated by one-way ANOVA followed by Dunnett’s test (****: p < 0.0001, blue asterisks). Statistical differences between the effects of the different β-arrestins were calculated using two-way ANOVA followed by Fisher’s test. (n.s.: non-significant; *: p < 0.05) *(For further details see Materials and Methods)* **(D)** Schematic cartoon depicting β-arrestins interacting with the GPCR in the tail-engaged conformation (as with the DRY mutant AT1R) and in the fully engaged form (with the wild-type AT1R). The method of BRET-based detection of the complex formation between the AT1R-Luciferase and β-arrestin1/2-mVenus constructs is also depicted. **(E-F)** BRET measurements of complex formation between the wild-type (WT – dark blue) or DRY-mutant (DRY – light blue) AT1R and β-arrestin1 **(E)** or the AT1R forms and β-arrestin2 **(F)** after stimulation with 100 nM AngII. The respective constructs were expressed in β-arrestin1/2 double KO HEK293 cells. Data are means ± SEM of four independent experiments, each performed in triplicates. **(G)** Bar diagrams showing the relative binding of β-arrestin isoforms to the WT or the DRY mutant AT1R. *Note the larger impact of the DRY mutation on β-arrestin2 binding to the AT1R.* Data were calculated from the ratios of AUC values of WT and DRY receptors in each separate experiment for both arrestins. Scatter plots display the individual AUC ratios. Statistical differences were evaluated to compare the WT/DRY AUC Ratios between β-arrestin1 and −2 using unpaired t-test (*: p < 0.05). **(H-K)** BRET measurements comparing the stability of complexes between AT1R and βarrestin1 **(H)**, or βarrestin2 **(I)** as well as between the AT1R and the AP2-binding mutant form of βarrestin1-AP2bm **(J)** or βarrestin2-AP2bm **(K)** after stimulation of 100 nM AngII followed by the application of 30 μM Losartan (AT1R antagonist) after 5 min. β-arrestin1/2 double KO HEK293 cells were transfected with AT1R-Luciferase and the mVenus tagged versions of β-arrestins or their AP2-binding mutants. Data are means ± SEM of four independent experiments, each performed in triplicates. **(L)** Bar diagrams showing the relative effects of the antagonist Losartan on the different AT1R-β-arrestin complexes calculated from the data shown in E-H. Values were obtained by calculating the ratios between the AUC values of DMSO vs. Losartan treated curves in each experiment for the time period when the antagonist was present. Data are means ± SEM of four independent experiments, each performed in triplicates. Statistical differences were calculated using two-way ANOVA followed by Fisher’s test. (n.s.: non-significant; *: p < 0.05)

These data together show that AP-2 binding is important to mitigate the ability of β-arrestin1 to desensitize AT1R while it had no impact on the more potent effect of β-arrestin2 on AT1R desensitization. Taken these data together with our previous observations, AP2 binding is important for β-arrestin1 to direct the AT1R to the CCP, and at the same time, AP2 binding of β-arrestin1 helps the receptor escape desensitization. Therefore, lastly, we wanted to determine how the two β-arrestins or their mutants interacted with the AT1R.

#### β-arrestin isoforms show structural differences in their interactions with AT1R

Stable GPCR-β-arrestin binding occurs via multiple molecular interactions between the two proteins and is also affected by the lipid composition of the surrounding membrane. Based on established views, stable interaction between the activated receptor and β-arrestin is achieved in consecutive steps. In the initial step, called tail-engagement, β-arrestins recognize the phosphorylated residues on the C-termini of the receptors. This interaction is followed by subsequent interaction between the so-called finger loop of β-arrestins and the cone-shaped cavity of the receptors formed between the transmembrane helices facing the intracellular aspect of the membrane ^74^. These two interactions establish the stable, fully engaged form of the receptor-β-arrestin complex (Fig. 6D). Since the alpha subunit of G-proteins binds to the same core-interface of the receptors marked by the highly conserved DRY residues ^75,76^ necessary for their activation, binding of β-arrestins to this site prevents G protein binding and, hence, is responsible for GPCR desensitization.

To investigate the relative contributions of the tail-engagement to the AT1R-β-arrestin interaction, we compared β-arrestin binding to either the wild-type AT1R or its DRY/AAY mutant, the latter being defective both in β-arrestin finger loop and G-protein interactions ^77,78^. The interaction between the arrestins and AT1Rs was analyzed in the β-arrestin1/2 knockout cells using BRET analysis, monitoring energy transfer between transfected mVenus-tagged β-arrestins and Luciferase-tagged AT1Rs. This analysis showed that, as expected, the DRY-mutant receptor had a reduced interaction with both β-arrestins compared to its wild-type form. Notably, however, this difference was significantly larger in the case of β-arrestin2 compared to β-arrestin1 (Fig. 6E-G). This suggested that β-arrestin2 relies upon binding to the core of the receptor via its finger loop more than β-arestin1. This finding was also consistent with the stronger ability of β-arrestin2 to uncouple the receptor from the G-proteins, as the interaction of the finger loop with the receptor is mutually exclusive with the receptor interaction with the G-protein alpha subunit.

To further assess the strength of AT1R-β-arrestin interactions, we tested the reversibility of the process using the non-peptide antagonist, losartan, added 5 min after Ang II stimulation. These experiments showed that β-arrestin2 dissociation from the receptors was slower than that of β-arrestin1 (Fig. 6H vs. Fig 6. I, the difference quantified in panel L). Notably, mutation of the AP2-binding residues in β-arrestin1 slowed down its dissociation from the receptors after Losartan addition, while the AP2-binding mutation did not significantly affect the already slower dissociation kinetics of β-arrestin2 (Fig. 6J and K, quantified in panel L).

All these data taken together were compatible with a model that posits that activated AT1R binds β-arrestin1, which carries the receptor to CCPs, where an interaction with AP2, promotes a partial release of β-arrestin1 from the receptor allowing a fraction of the receptors to reengage with G-proteins. In contrast, β-arrestin2, which is already prepositioned in CCPs and shows stronger binding to the receptors partly provided by its finger-loops, keeps the receptors in a desensitized state and more efficiently directs them for internalization.

## DISCUSSION

In the present study we focused on the process of GPCR re-sensitization, a poorly understood aspect of GPCR function. Many GPCRs show rapid desensitization, i.e. uncoupling from their cognate G-protein partner, and at the same time undergo endocytosis, mostly via clathrin-mediated endocytosis (CME). Ligand-induced endocytosis also decreases the available receptors on the surface, a process termed down-regulation ^5^. According to current views, desensitized GPCRs must be internalized and undergo a series of trafficking steps to reappear in the PM and regain their G protein activating competence. Central players in all of these processes are β-arrestins, whose binding to the activated receptors is responsible for both desensitization and internalization ^79^. An important aspect of these molecular events is the phosphoinositide content of the PM, more specifically, PI(4,5)*P*_2_, which is critical for the association of many components of CME ^80–82^ but is also involved in GPCR - β-arrestin interactions ^33,37,38,65^}.

The present studies highlight the importance of PM PI(4,5)P_2_ in GPCR re-sensitization and point to the critical roles of PIP5K1A and EFR3A in the process. As for EFR3, it was the Rbo mutation in Efr3 in *Drosophila* that showed rapid loss of light response in the eye ^83^, and was also linked to endocytosis ^84^. Our earlier studies showed that EFR3s were also important for AT1R re-sensitization ^44^, which is now attributed to EFR3A based on the current results. Moreover, the present study also showed the PIP5K1A, specifically, is also required for AT1Rs to maintain their G protein coupling. An early study from the Lefkowitz group reported the role of PIP5K1A in the production of the PM PI(4,5)*P*_2_ pool used for CCP assembly and therefore, promoting the sequestration of β-Adrenergic receptor in CCPs ^85^. Another work showed that PIP5K1A plays an important role in CCP initiation and maturation ^39^. A more recent study found that β-arrestin2 recruits PIP5K1B to the PM to regenerate the PM PI(4,5)*P*_2_ pools after PLCβ activation by protease activated receptors 2 ^86^.

An important conclusion of the current study was that while EFRs and PIP5K1A were, indeed, important for AT1R endocytosis, a full endocytosis cycle was not necessary for at least a fraction of the receptors to maintain their G protein coupling. Instead, we found that sorting the receptors to CCPs and the presence of the tetrameric adaptor, AP2, was critical for AT1R re-sensitization. The importance of PI(4,5)P_2_ in AP2 binding to the PM has been well documented ^81,87,88^ and as pointed out earlier, PIP5K1A was found to play a particularly important role in CCP maturation ^39^. Since all these factors are also important for endocytosis, previous studies logically reached the conclusion that endocytosis is a prerequisite for GPCRs to re-sensitize.

An unexpected finding of the current studies was that the two β-arrestins showed clear differences in their roles in the process of AT1R re-sensitization. While β-arrestin1 binding to the AT1R protected the receptor from fast desensitization, β-arrestin2 strongly promoted the uncoupling of the receptor form the Gq protein and directed it to endocytosis. This prompted us to explore differences between the two β-arrestins in their relationship to CCPs, AP2 binding and interaction with the receptor. These studies showed that β-arrestin2 already showed association with CCPs even before receptor activation, and AP2 knock-down or mutating its AP2 binding site had relatively little effect of its recruitment to CCPs. In contrast, β-arrestin1 binding to CCPs required receptor activation and its recruitment to CCPs was sensitive to AP2 knockdown or mutating its AP2 binding site. Additionally, the two β-arrestins showed important differences in their interaction with the activated AT1Rs. The interaction of β-arrestin2 with the receptor relied more on the finger-loop binding to the receptor core than in the case of β-arrestin1, which explains why the former interferes with G protein-coupling more effectively. Curiously, mutations of β-arrestin1 AP2 binding site strengthened its interaction with the AT1R and its ability to desensitize the receptors when cells lack β-arrestin2. These data suggest that β-arrestin1 takes a fraction of AT1Rs to CCPs where β-arrestin1 interaction with AP2 facilitates the release of the receptor from β-arrestin1 and allow its new round of G protein activation. In contrast, the stronger binding of β-arrestin2 to the receptor and its lesser reliance on AP2 interaction, makes it more potent to desensitize the receptors and direct them for internalization. Importantly, increasing number of studies reported differences between β-arrestin1 and β-arrestin2 in GPCR signaling, internalization and desensitization ^89–91^, including the limited desensitizing effect of β-arrestin1^92^.

The importance of PI(4,5)P_2_ and the roles of PIP5K1A and EFR3 in this process is likely to be two-fold: both via directly affecting the PM binding and receptor interaction of β-arrestins or indirectly, through interference with AP2 function. Direct PI(4,5)*P*_2_ binding of β-arrestin isoforms was revealed and confirmed in several recent structural studies ^37,38,93^ and its functional role was also explored. Those studies concluded that PI(4,5)*P*_2_ is important for stabilizing GPCR-β-arrestin complexes, and determining the fate of this complex ^65^. This latter elegant study has analyzed the PI(4,5)*P*_2_ dependence of the interaction of the two β-arrestins with a large panel of GPCRs using mutant β-arrestins that are impaired in PI(4,5)*P*_2_ interaction. Their analysis found relatively little difference between the two β-arrestins in their lipid dependence, but in general, β-arrestin1 showed a higher lipid dependence than β-arestin2, which was more substantial with some receptors, such as the oxytocin receptor, and even reversed in the case of tachykinin receptors ^65^. This study has uncovered a large difference between receptors whose β-arrestin interaction showed strong, while other displayed minimal PI(4,5)*P*_2_ dependence. Importantly, according to that study, AT1R occupied the area between these two groups of receptors. In our studies, using the AT1R, we found that depletion of the levels of EFR3A or PIP5K1A caused a clear shift, favoring β-arrestin2 binding to the receptor on the expense of β-arrestin1 and these changes were associated with a faster GPCR desensitization. It is also worth pointing out that, in our studies, knock-down of either EFR3 or PIP5K1A caused faster desensitization of both endogenous β-adrenergic receptors and V2 vasopressin receptors, even though they were found vastly different in their PI(4,5)*P*_2_ dependence of their β-arrestin recruitment in the Janetzko study.

As for PI(4,5)*P*_2_ dependence of its binding, the best characterized component of CCPs is the adaptor protein AP2 ^87,88^. This tetrameric adaptor has several of its subunits showing strong interaction with the lipid each of which controlling distinct functions of the complex ^94^. Previous observations ^39^ and our current results suggest that a specific pool of PI(4,5)P_2_ generated by an EFR3 and PIP5K1A dependent process controls AP2 interaction with the PM and the β-arrestin GPCR complex. It is most likely, that β-arrestins assume different conformations when they interact with different GPCRs (see ^37,38,93^, which will impact not only their PI(4,5)P_2_ dependence (as shown in ^65^), but also their ability to interact with AP2 and interfere with the G protein coupling of the receptors. It has been shown that in some cases, G protein activation is still possible while the receptor still binds β-arrestin1 and the existence of GPCR-β-arrestin-G-protein megacomplexes have been already reported ^95,96^. The conformational flexibility of β-arrestins, together with differences between its two forms adds to the fine-tuning of the unique signaling modality of the various GPCRs.

In summary, our studies show that the ability of AT1Rs to maintain their G protein signaling competence requires their sorting into AP2 positive CCPs but not their internalization. This process depends on a PM PI(4,5)*P*_2_ pool that is specifically produced by PIP5K1 with the involvement of EFR3A, which also determines the balance between β-arrestin1 and −2 interaction with the receptor, the former protecting it from rapid desensitization mostly mediated by β-arrestin2. While desensitization of the tested β1-adrenergic and V2 receptors was similarly enhanced by depletion of PIP5K1 and EFR3, it is most likely that significant variations exist between GPCRs in their β-arrestin interactions and their dependence on PM PI(4,5)*P*_2_ as they relate to receptor desensitization and re-sensitization.

## Key resources table

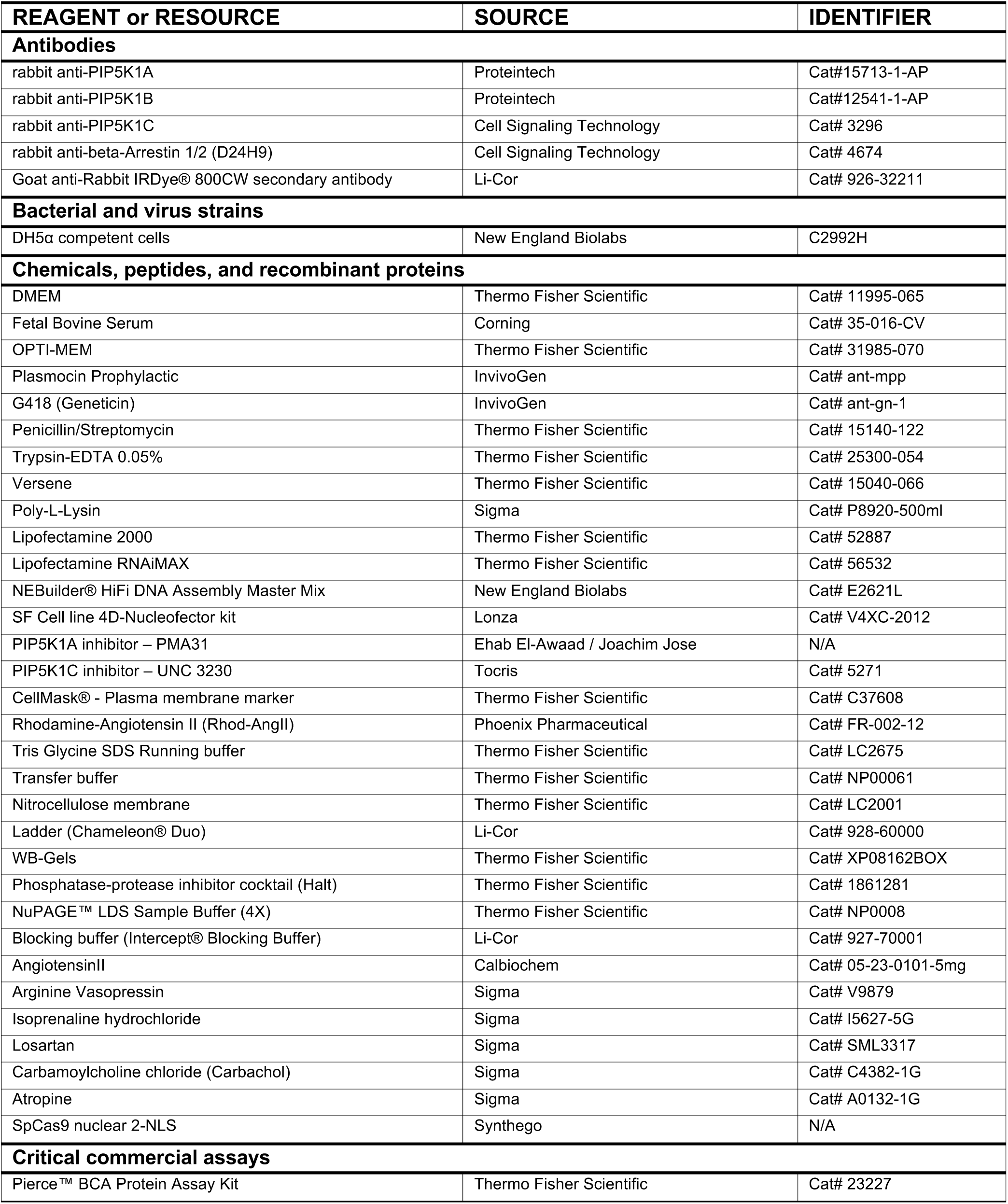
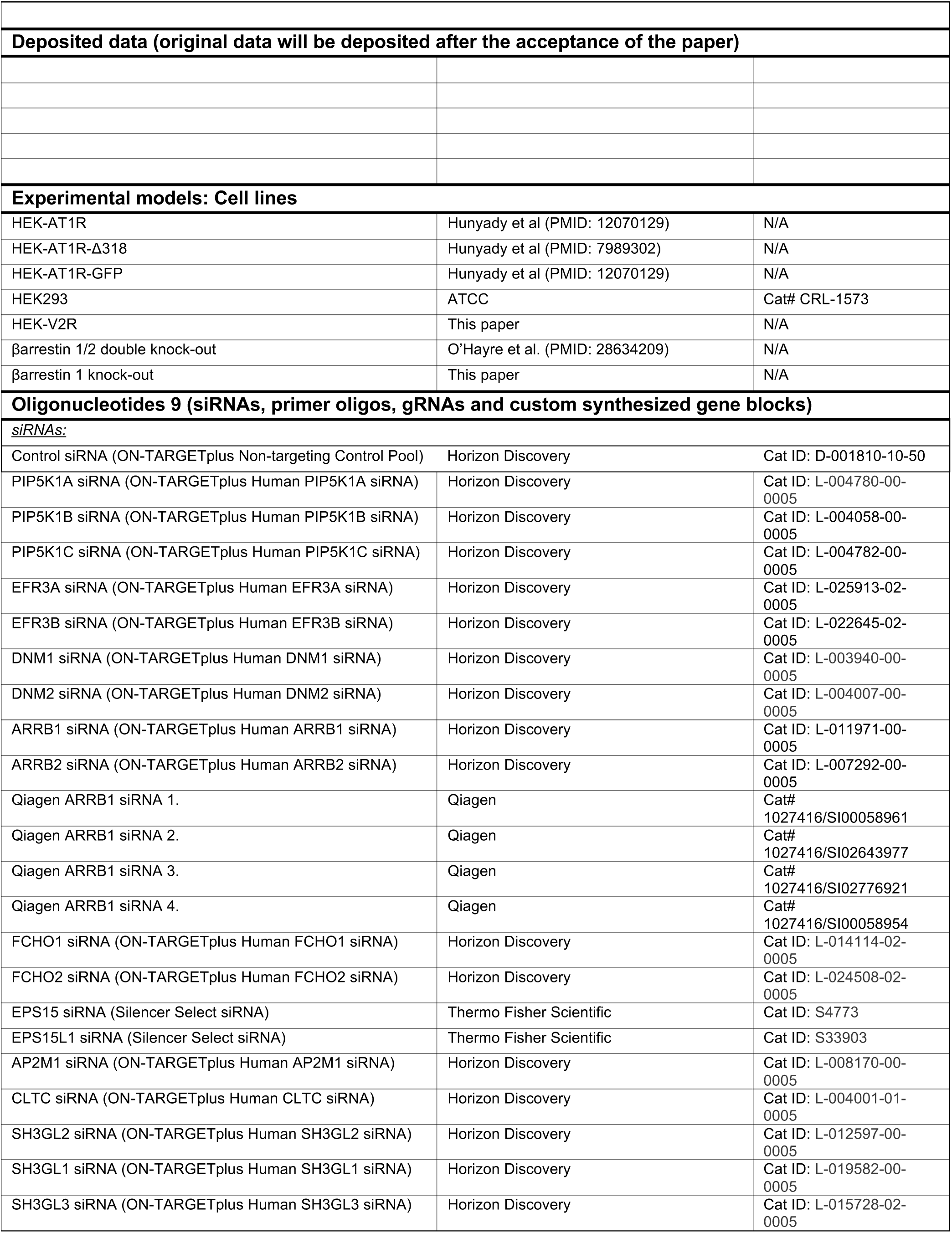
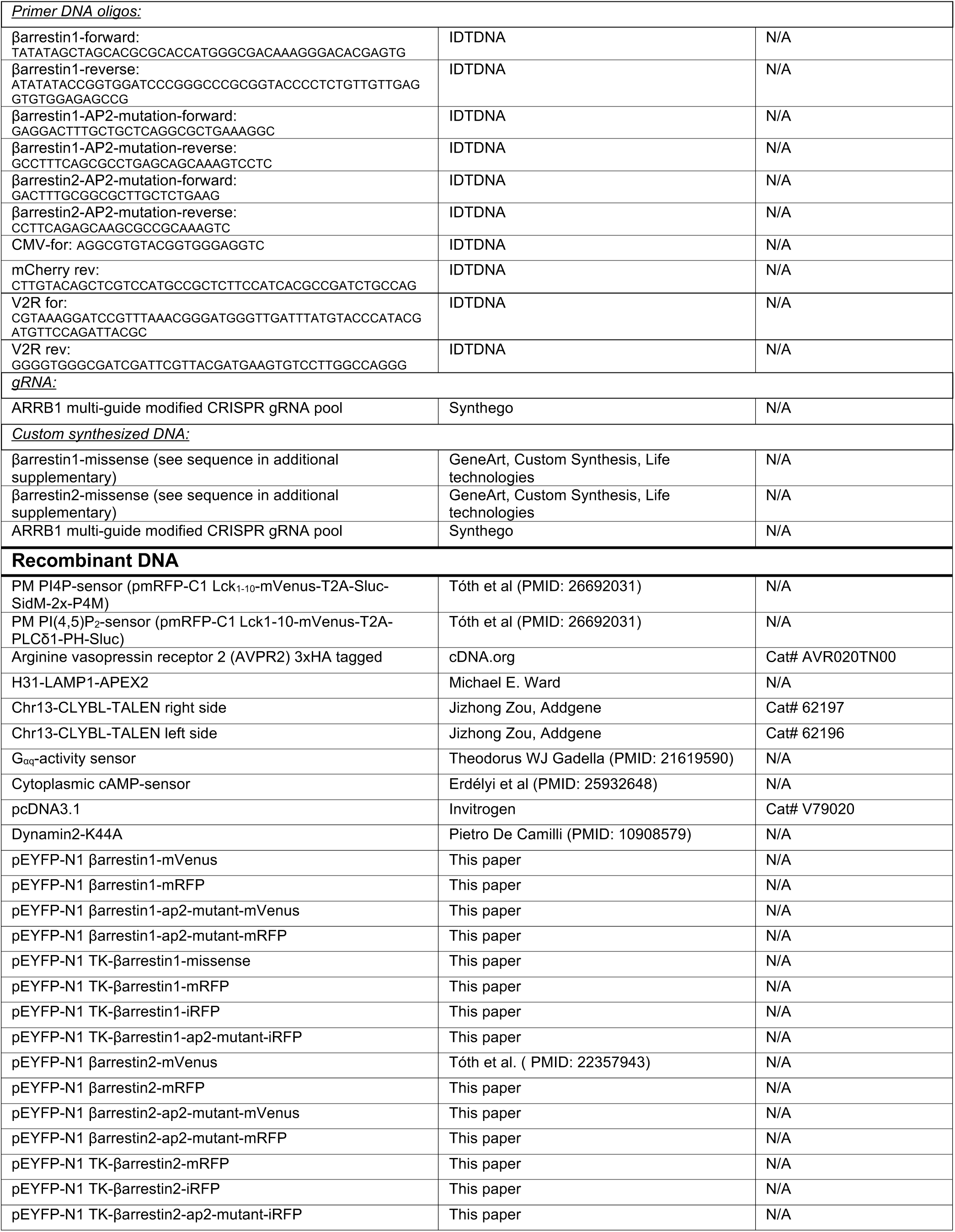
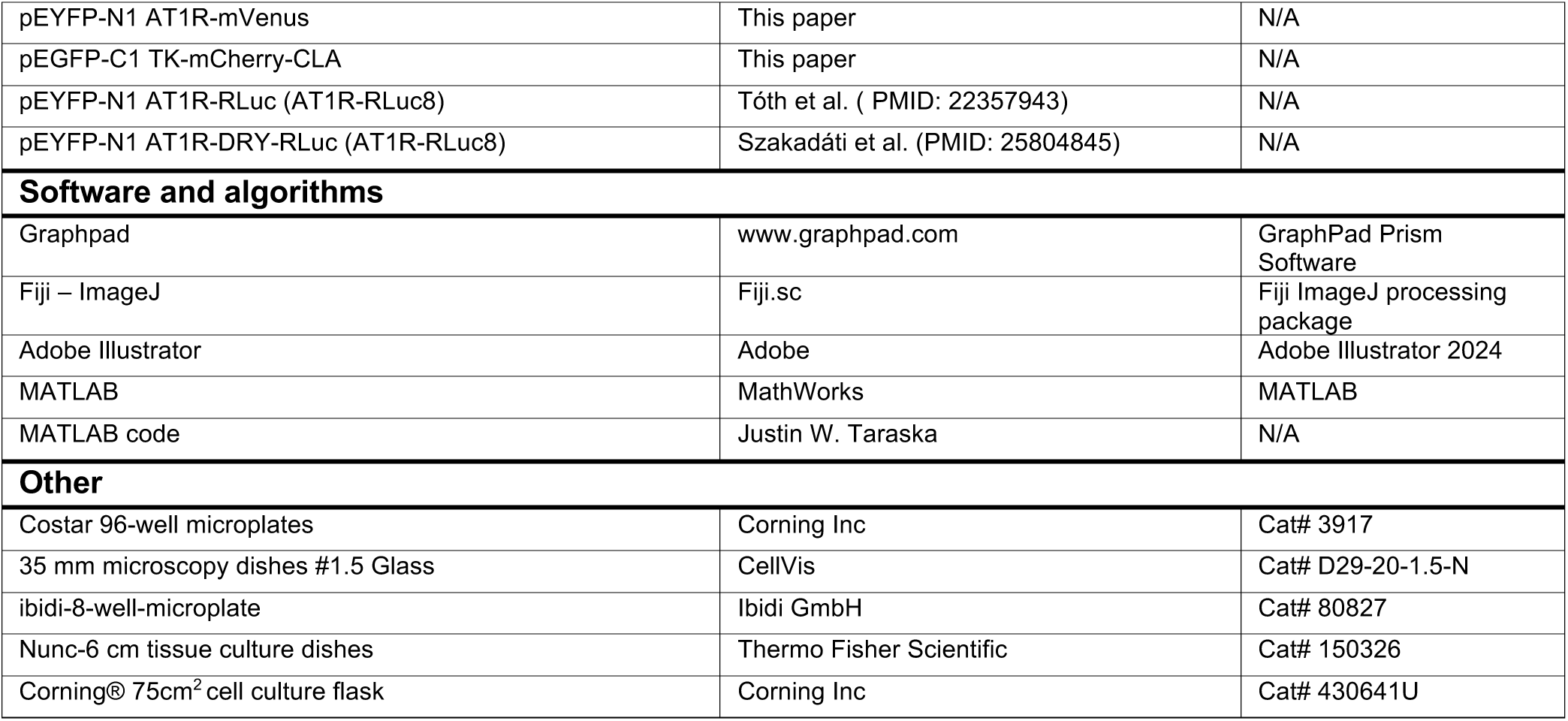

### RESOURCE AVAILABILITY

#### Lead contact

Further information and requests for resources and reagents should be directed to and will be fulfilled by the lead contact, Tamas Balla.

#### Materials availability

Plasmids generated in this study are available from the lead contact upon request without restrictions.

#### Data and code availability

∘ Original western blot images have been deposited at Mendeley and are publicly available as of the date of publication. The DOI is listed in the key resources table.
∘ Microscopy data reported in this paper will be shared by the lead contact upon request.
∘ Any additional information required to reanalyze the data reported in this paper is available from the lead contact upon request.

### EXPERIMENTAL MODEL AND STUDY PARTICIPANT DETAILS

Cell cultures used in this study were maintained in 10% fetal bovine serum supplemented Dulbecco’s Modified Eagle’s Medium (DMEM) also containing 1% penicillin/streptomycin and prophylactic (5μg/ml) Plasmocin. Cells were regularly tested for mycoplasma contamination. Cells were kept in 5% CO2 at 37°C humidified incubator on 75 cm^2^ cell culture flasks.

### METHOD DETAILS

#### Materials

All materials used in this study are described in the Key Resource Table. Restriction enzymes and T4 DNA ligase were obtained from New England Biolabs.

#### DNA constructs

In this paper, rat β-arrestin sequences were used. The β-arrestin2-mRFP construct was created by replacing mVenus sequence in β-arrestin2-mVenus using AgeI and NotI restriction sites. β-arrestin1-mVenus and -mRFP constructs were generated by replacing the sequence of β-arrestin2 with the amplified sequence of β-arrestin1 using the NheI and AgeI sites. For the PCRs, β-arrestin1-GFP was used as template (kindly provided by Marc G. Caron). TK-promoter driven versions were made by replacing the promoter sequences with the sequence of the thymidine-kinase promoter from the TK-YFP-STIM1 construct ^97^ using AseI and NheI sites. TK-β-arrestin1-iRFP and TK-β-arrestin2-iRFP constructs were generated by replacing mRFP sequence with the sequence of iRFP (pmiRFP-N1, ^98^) using AgeI and NotI restriction sites. AP2 binding mutations (R393,395A for β-arrestin1; R394A/R396A for β-arrestin2) were introduced using site directed mutagenesis. The siRNA-resistant form of β-arrestin1 sequence was obtained from Life Technologies as GeneArt Custom Synthesis (for the sequence, see the Supplementary data). The TK-promoter driven siRNA-resistant β-arrestin1 construct was generated by replacing the β-arrestin1-mRFP sequence in the TK-β-arrestin1-mRFP construct with the siRNA-resistant β-arrestin1 sequence using the NheI and NotI sites. AT1R-mVenus was made by replacing Rluc8 sequence with the sequence of mVenus (amplified from Lck_1-10_-mVenus, ^99^). The pEGFP-C3 EGFP-CLA-light chain construct was a kind gift from Lois Greene (NHLBI). The TK-GFP-CLA construct was made by replacing the CMV promoter sequence with the thymidine-kinase promoter obtained from TK-YFP-STIM1 construct using the AseI and NheI sites. The TK-mCherry-CLA construct was prepared from the EGFP-tagged version by replacing the GFP sequence with the mCherry sequence amplified from pmCherry-using AgeI and BglII digestion and ligation. All constructs were verified by sequencing.

#### Western Blotting

For Western Blots of experiments where siRNA knock-down was used, cells were plated at a density of 5×10^4^ cells/well in 6-well plates in culture medium at 37°C (Day 1). One day later, cells were transfected with 200 µl transfection mixture containing 100 pmol specific siRNA, and 5 µl/dish Lipofectamine RNAiMAX (Day 2). After 20 hours the media was replaced with fresh culture medium (Day 3). On the following day (Day 4), cells were transfected again with 200 µl transfection mixture containing the indicated DNA constructs, 2 µl/dish Lipofectamine 2000, and after 4-6 hours the culture media was replaced. On Day 5, cell culture plates were placed on ice, cells were rinsed with ice-cold PBS and scraped into 200 μl RIPA buffer (also containing 1000x phosphatase and protease cocktail (Thermo Fisher Scientific). After centrifuging the samples at 15,000 x g, (10 min, 4 °C), the protein concentration was calculated by a BCA assay. Aliquots containing equal amounts of protein were complemented with 4x-LDS sample buffer and boiled at 95 °C for 5 min before resolving on 8-16 % SDS gels by running at 230 V for 90 min at R/T. Proteins were transferred to nitrocellulose membranes and incubated with the respective primary (1:1000) and secondary antibodies (1:5000). Membranes were blocked at R/T for 1 hour using the LI-COR Blocking Buffer before applying primary antibodies and rinsed for 5 minutes three times in PBST before and after incubations with the secondary antibody. The blots were then scanned by an Odyssey fluorescence imager (LI-COR, NE). In experiments assessing the selected knock-out cell clones, the Western Blot procedure was simplified. After the second passage of the clonal cultures, 1×10^6^ cells were centrifuged and resuspended directly in 200 μl RIPA buffer (containing 1000x phosphatase and protease cocktail) and processed further as described above.

#### Bioluminescence Resonance Energy Transfer (BRET) measurements

For BRET measurements, 1.2×10^4^ cells/well were seeded onto poly-L-lysine-coated white 96-well, flat bottom microplates (Corning). Poly-L-Lysine (Sigma, P8920, diluted to 0.001%) was added to the wells for 1 h, and after removal let the plates dry before cell plating. In the siRNA experiments, one day after seeding (Day 2) cells were transfected with the specific siRNA constructs (50 µl/well transfection mixture containing 15 pmol siRNA, and 0.75 µl/well Lipofectamine RNAiMAX in OPTI-MEM reduced serum transfection media. 20 hours later (Day 3) the culture media was replaced with fresh culture medium (DMEM, 10% FBS, 1% Pen/Strep). On the 4^th^ day, cells were transfected again with the indicated DNA constructs in 50 µl transfection mixture containing the indicated DNA constructs (25-50 ng/well for CMV-promoter driven plasmids and 300-400 ng/well for TK-promoter driven plasmids), and 0.5 µl/dish Lipofectamine 2000 and the media was replaced 4-6 hours later. BRET measurements were performed 25-26 hours after transfection (this time window was critical). For BRET measurements, the cell culture media was replaced by 50 µl assay-media (modified Krebs-Ringer buffer containing 120 mM NaCl, 4.7 mM KCl, 1.2 mM CaCl_2_, 0.7 mM MgSO_4_, 10 mM glucose, and 10 mM Na-HEPES, pH 7.4) and incubated for 30 mins at 37°C. The cell permeable luciferase substrate, Coelenterazine h, was freshly dissolved and added to the cells in 40 µl/well volume in the same buffer to yield a final concentration of 5 µM. For the measurements, Tristar2 LB 942 Multimode Microplate Readers (Berthold Technologies) was used at 37°C. Intensities were recorded for 500 ms/well using 475/20 and 540/40 nm emission filters. After recording a baseline, the indicated reagents, dissolved in modified Krebs–Ringer buffer, were added in 10 µl volume with manual pipetting. For this, the plates were unloaded resulting in the interruption of the recordings. To calculate the BRET ration changes for plasma membrane lipid or the cytoplasmic cAMP measurements, a previously published workflow was followed ^100^. To calculate the changes in the association between the AT1R-Luciferase and mVenus-tagged β-arrestin constructs, first the BRET ratio values were calculated by dividing the values of AngII-treated cells with those of the DMSO treated ones. To assess the extent of increase, these curves were then normalized to the average of the last 5 points before the stimulation.

#### Fluorescence Resonance Energy Transfer (FRET) measurements

For FRET measurements, HEK-AT1R cells were plated on poly-L-lysine-treated glass-bottom IBIDI 8-well μSlides at 1.5×10^4^ cells/well density. One day after seeding, cells were transfected with the indicated siRNAs, (at 20 μl/well transfection mixture containing 20 pmol/well siRNA and 0.75 μl/well Lipofectamine RNAiMAX transfection reagent dissolved in OPTI-MEM) following the manufacturer’s recommendations. After 20 hours, the cell culture media was replaced and one day later, the cells were transfected with the FRET-based Gq-activity sensor ^55^ (200 μl transfection mixture containing 0.5 μl/well Lipofectamine 2000 and 200 ng of the sensor’s DNA construct dissolved in OPTI-MEM). After 4-6 hours, the cell culture media was replaced with fresh culture media and FRET measurements were performed 20-26 hours after transfection. For this, the medium was changed to 300 µl assay-media (modified Krebs-Ringer buffer containing 120 mM NaCl, 4.7 mM KCl, 1.2 mM CaCl_2_, 0.7 mM MgSO_4_, 10 mM glucose, and 10 mM Na-HEPES, pH 7.4) and the culture dish was mounted to the stage of an Olympus IX73 inverted microscope equipped with a Lambda 421 illuminator (Sutter Instruments) and a Hamamatsu ORCA Flash 4.0 CMOS camera. A Hamamatsu W-View Gemini system with a 500 nm dichroic mirror was used to split donor (475 nm) and acceptor (525 nm) wavelengths to the two halves of the camera sensor and recordings were made at RT. For data acquisition and analysis, the MetaFlour (Molecular Devices) software was used. The indicated reagents were dissolved in modified Krebs–Ringer buffer and were added manually in 100 μl volume. The simple acceptor to donor ratios were formed by dividing the mVenus fluorescence intensities with those of the mTurqoise2 intensities and normalized to their pre-stimulatory values.

#### Internalization assay by confocal microscopy

HEK293-AT1 cells were seeded at a density of 1.5×10^4^ cells/well on poly-L-lysine-treated glass bottom IBIDI 8 well μSlides and cultured at 37°C. For siRNA transfection, one day after seeding the culture medium was replaced with 200 µl transfection mixture containing 20 pmol/well siRNA and 0.75 μl/well Lipofectamine RNAiMAX transfection reagent dissolved in OPTI-MEM. 20 hours later the media was replaced with fresh culture media and the cells were kept in a 5% CO_2_, 37°C incubator for two more days. Before the experiments, the media was replaced with 300 µl assay-media (modified Krebs-Ringer buffer containing 120 mM NaCl, 4.7 mM KCl, 1.2 mM CaCl_2_, 0.7 mM MgSO_4_, 10 mM glucose, and 10 mM Na-HEPES, pH 7.4) containing CellMask® at 3000x dilution (to mark the plasma membrane) and 30 nM Rhodamine-tagged angiotensin II (Rhod-AngII). The plates were kept in dark at 37°C (no CO_2_) for 20 minutes. After the incubation period, cells were placed on ice and rinsed with ice-cold PBS twice before fixing with 4%-PFA solution. Three images from different areas were taken from every dish using a Zeiss LSM 710 confocal microscope equipped with a 1.4 NA, 63x objective. For calculating the cytoplasmic/PM intensity Ratios of Rhod-AngII, images were background subtracted, and a binary mask was created to mark the plasma membrane. and cytoplasmic intensities based on the contour of the PM-staining CellMask signal. Using this mask, intensities associated with the PM and the cytoplasmic were calculated and divided to generate the cytoplasmic/PM-ratio. Post-acquisition picture analysis was performed using Fiji.

#### TIRF microscopy

HEK-AT1R or HEK-AT1R-GFP cells were plated on poly-L-lysine-treated glass bottom IBIDI 8-well μSlides at 1.5×10^4^ cells/well density. On the following day, cells were treated with the indicated siRNAs (15 pmol/well) dissolved in 200 μl OPTI-MEM also containing 0.75 μl Lipofectamine RNAiMAX). After 20-24 hours the media were replaced with fresh culture media. The following day, cells were transfected with the indicated TK-β-arrestin construct (200 ng/well) in 200 μl OPTI-MEM containing 0.5 μl Lipofectamine 2000 per well. The media was changed after 4-6 hours and cells were analyzed 20-24 hours later using a Nikon Eclipse TiE inverted microscope equipped with an iLas2 (Nikon) TIRF module and a 1.45 NA, 100x TIRF objective. Fluorescence images were captured by an ORCA-Fusion BT digital CMOS camera (Hamamatsu). For the measurements, cells were kept at 30°C on a heated stage and heated objective in 300 µl assay-media (modified Krebs-Ringer buffer containing 120 mM NaCl, 4.7 mM KCl, 1.2 mM CaCl_2_, 0.7 mM MgSO_4_, 10 mM glucose, and 10 mM Na-HEPES, pH 7.4). After a 40 seconds of control period, cells were stimulated with AngII (100 nM final, added manually in 100 μl assay media). Images were acquired in every 5 seconds. To calculate the changes in normalized fluorescence intensities, images were background subtracted, then the average intensities were estimated for every cell using ROIs and excluding the pixel values of 0. For normalization, each average intensity value was divided by the average pre-stimulatory value calculated from the last three time points before the stimulation. Post-acquisition picture analysis was performed using Fiji software.

#### Fluorescent Lifetime Imaging Microscopy

HEK293 cells with β-arrestin 1/2 double knock-out were seeded on 35 mm dishes at a density of 3×10^5^ cells/dish. On the following day, cells were transfected with 200 μl transfection mixture containing the Gq-activity sensor (1 μg/dish) and the indicated β-arrestin construct (300 ng/dish) together with 2 μl/dish Lipofectamine 2000. Culture media was replaced 4-6 hours later, and cells were analyzed 20–24 h post transfection in a modified Krebs-Ringer buffer (containing 120 mM NaCl, 4.7 mM KCl, 1.2 mM CaCl_2_, 0.7 mM MgSO_4_, 10 mM glucose, and 10 mM Na-HEPES, pH 7.4) at room temperature using an Olympus IX81 inverted microscope. The fluorescent lifetime of mTurqoise2 was calculated based on the frequency-domain from images recorded in 12 phases with a cooled Lambert Toggel – MDL1.1camera and a pulsed LED laser illuminator LED (L6CC-CSB Oxxius Simply Light, all assembled by Lambert Instruments). On the day of the experiments a reference lifetime value was recorded using a Fluorescein standard (Lambert Instrument, lifetime: 4.02 ns). Images were acquired in every 10 seconds. 100 nM AngII was dissolved in 1 ml assay-media and added after four control acquisitions. Fluorescent lifetimes were estimated for each individual cell using ROIs and calculating the average lifetime based the frequency domain. To calculate fold-changes, every life-time value was divided by the average lifetime values recorded for the control period. For image acquisition and analysis, the LIFA software was used.

#### Experiments assessing association with Clathrin-coated pits

Β-arrestin 1/2 double knock-out HEK293 cells were plated on poly-L-lysine-treated glass bottom IBIDI 8-well μSlides at 5×10^4^ cells/well density. One the following day, cells were transfected with the indicated DNA constructs by replacing the culture media with 200 μl transfection mixture containing 10 ng/well AT1R-mVenus, 300 ng/well TK-mCherry-CLA and 400 ng/well TK-β-arrestin-iRFP DNA and 0.5 μl Lipofectamine 2000 reagent. Four to six hours after transfection the culture media was replaced and cells were imaged after 20–24 h after transfection in 300 μl modified Krebs-Ringer buffer (containing 120 mM NaCl, 4.7 mM KCl, 1.2 mM CaCl_2_, 0.7 mM MgSO_4_, 10 mM glucose, and 10 mM Na-HEPES, pH 7.4) at 30°C on a heated stage and heated objective (1.45 NA, 100x) using the same TIRF microscope described above. Images were taken in every 5 seconds for 10 minutes. For correlative image analysis, selected frames were chosen (before stimulation and 1, 5 or 9 minutes after stimulation with 100 nM AngII). The recorded channels were split, and images of the individual channels were background subtracted, and arranged as β-arrestin-clathrin or AT1R-clathrin image pairs. These images were then analyzed for co-localization using a MATLAB code kindly provided by Justin W. Taraska ^70^. In this analysis, the clathrin channel is segmented based on every clathrin-coated structure and the correlation was calculated between the related other channel within a 6 nm radius circle around the center (local maxima) in each segmented clathrin structure.

#### Generation of the HEK293-V2R cell line

The vasopressin 2 receptor (V2R) stably expressing cell line (HEK293-V2R) was generated by targeting the CLYBL safe-harbor region on chromosome 13 by a TALEN based method and inserting the human V2R sequence after a tetracycline inducible promoter sequence. This system was described in detail in Cerbini et al. ^101^ and its modified version containing the the LAMP1-APEX2 sequence was kindly provided by Dr. Michael E. Ward ^102^. Using this plasmid, the LAMP1-APEX2 sequence was replaced between the PmeI and NruI sites by the amplified human V2R sequence using Gibson assembly (NEBuilder® HiFi DNA Assembly Master Mix, NEB). For targeted gene delivery, HEK293 cell were plated in 5×10^5^ cells/well density on poly-L-lysine-treated 6-well plates in culture medium at 37°C and the following day were treated with 200 μl transfection mixture containing 1 μg each of the DNA constructs (left and right TALEN plasmids and the plasmid encoding the V2R sequence) and 5 μl Lipofectamine LTX reagent. 6 hours later, the culture media was replaced and from the following day, the cells were treated with 500 μg/ml G418 for 7 days. Single cell clones were then grown in 96 well plates and amplified. Positive clones were selected based on their cAMP response to 1μΜ AVP stimulation in BRET experiments.

#### Generation of the β-arrestin1 knock-out cell line

β-arrestin1 knock-out cells were generated by CRISPR/Cas9. Commercially available synthetic gRNA pools (3 gRNAs per pool) targeting the ARRB1 gene was obtained from Synthego, along with recombinant SpCas9 enzymes. gRNAs and the Cas9 enzyme was then incubated in Eppendorf tubes for 30 minutes in SF Cell line 4D-Nucleofactor solution (Amaxa). HEK-AT1R cells were trypsinized and resuspended in the same SF Cell line 4D-Nucleofactor solution at a 6×10^5^ cells/100 μl density and mixed with the incubated ribonucleoproteins (RNP). Cells were then subjected to nucleofection using the AMAXA 4D-Nucleofactor (program: CM-130) following plating on 6 cm cell culture dishes. After three days, cells were trypsinized and single clone plated on 96 well plates. After clonal progression, cell clones were subjected to functional testing in BRET experiments by measuring plasma membrane PIP_2_ responses to AngII and measuring their β-arrestin1 and −2 expression levels by Western Blotting. Selected clones were then subjected to genomic sequencing and analysis using the ICE CRISPR Analysis Tool (Synthego).

#### Statistical analysis, data plotting

For data plotting and statistical analysis, the GraphPad Prism (San Diego, CA) software was used. For the specific statistical analyses see the Figure legends for each experiment.

## ACKNOWLEDGEMENT

We are grateful to Drs. Ehab El-Awaad and Joachim Jose (University, Munster, Germany) for the PIP5KA inhibitor, PMA-31. We are grateful for Dr. Justin Taraska (NHLBI, NIH) for sharing the MATLAB code assessing co-localization. This work was partly supported by the Intramural Research Program of the *Eunice Kennedy Shriver* National Institute of Child Health and Human Development of the National Institutes of Health. Confocal imaging was performed in the Microscopy Core of NICHD with the kind assistance of Drs Vincent Schram and Ling Yi.

## LEGENDS TO SUPPLEMENTARY FIGURES

**Figure S1.**
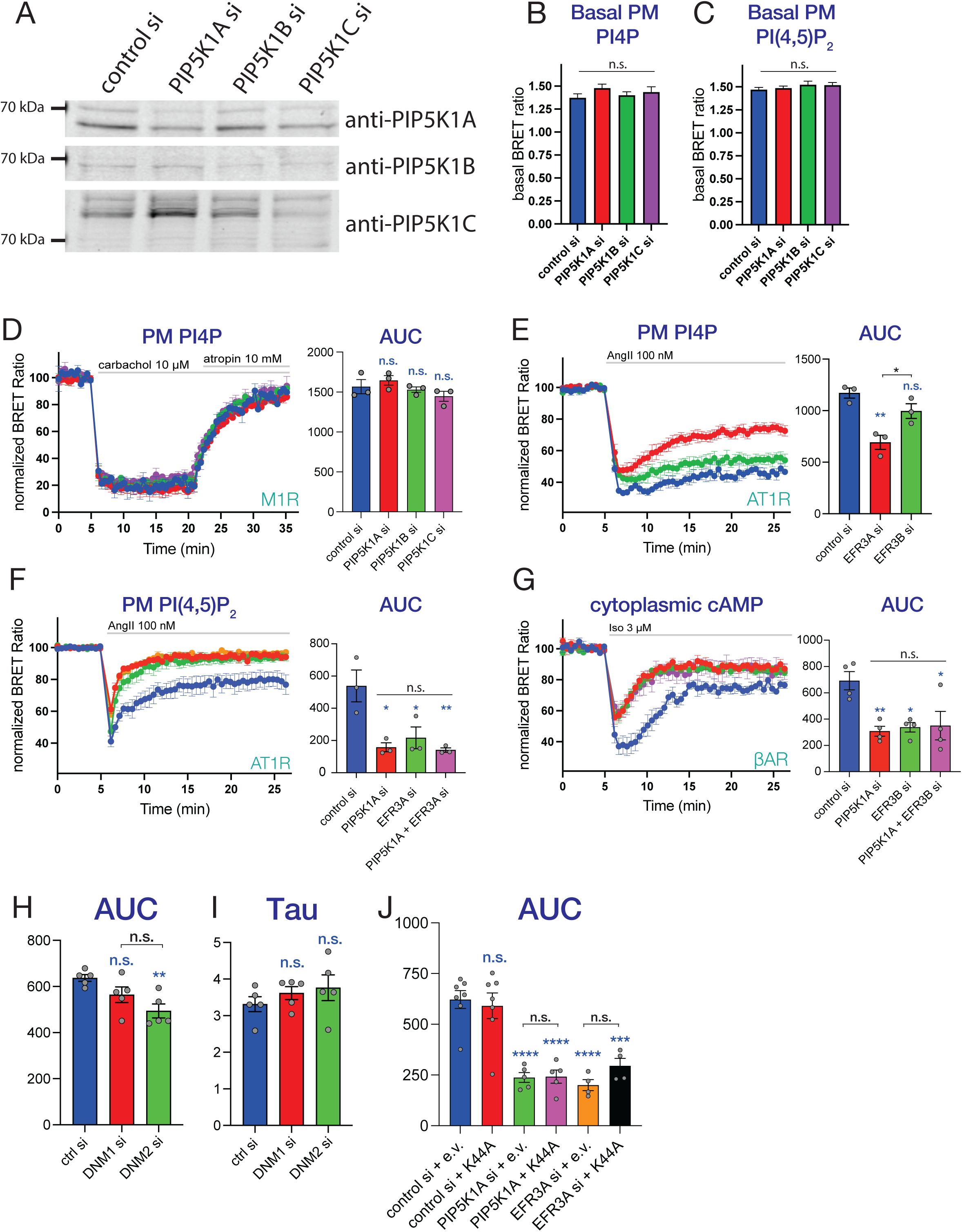
Monitoring PM PPIns levels in resting and stimulated cells after knocking-down distinct PIP5K1A and EFR3 isoforms. (**A**) Representative Western Blot analysis of HEK-AT1 lysates from cells treated with the indicated isoform specific siRNA against PIP5K1 proteins. (**B**) PM PI4P and **(C)** PI(4,5)P_2_ levels in unstimulated HEK-AT1 cells after silencing of distinct PIP5K1 isoforms. Basal PM PPIns levels were assessed by the resting BRET-Ratio values obtained before stimulation in each experiment in cells expressing the PI4P and PI(4,5)P_2_ BRET sensors. Data are means ± SEM of three **(B)** or four **(C)** independent experiments, each performed in triplicates. Statistical difference was calculated using one-way ANOVA followed by Tukey’s post-hoc test. (n.s: not significant) **(D)** Monitoring changes in PM PI4P levels after M1R activation in HEK-AT1 cells knocked down for the indicated PIP5K1A isoforms (control – blue, PIP5K1A – red, PIP5K1B – green, PIP5K1C – purple) using a BRET analysis. After a 5 min control period, cells were treated with 100 μM carbachol (CCh) followed by the addition of 10 μM atropine after 15 mins. Bar graphs show area under the curve (AUC) calculations for the BRET values covering the recovery (atropine) period to evaluate statistical differences between the groups. Data are means ± SEM of three independent experiments, each performed in triplicates. Scatter plots show results of individual experiments. Statistical significance was obtained by using one-way ANOVA followed by Tukey post-hoc test to estimate differences between the separate groups in multiple comparisons. (n.s: not significant) **(E)** Monitoring changes in PM PI4P levels after AT1R activation in HEK-AT1 cells treated with the indicated targeting siRNAs. Cells were stimulated with 100 nM AngII after a 5 min control period. Data are means ± SEM of three independent experiments, each performed in triplicates. Columns show mean ± SEM of AUC calculations scatters representing the individual values. One-way ANOVA followed by Tukey’s post-hoc test was used for evaluation of the statistical differences in multiple comparisons. Blue asterisks display the differences compared to the control, while black asterisks represent differences between the different treatment groups (n.s.: non-significant; *: p<0.05; **: p < 0.01). **(F-G)** Comparisons of single or double knockdown of PIP5K1A and EFR3 isoforms on the activity of **(E)** AT1R or **(F)** β-adrenergic receptors (βΑR) using BRET-based measurement of PM PI(4,5)P_2_ or cytoplasmic cAMP levels, respectively, in HEK-AT1R cells treated with the indicated siRNA(s). Cells were stimulated with 100 nM AngII to stimulate AT1R **(F)** or with 3μM isoprenaline (Iso) **(F)** to stimulate endogenously βARs after a 5 min control period. Bar graphs show area under the curve (AUC) calculations for the curves in both experiments to compare the separate groups. Scatter plots show the results of individual experiments. Data are means ± SEM of **(F)** three or **(G)** four independent experiments, each performed in triplicates. Statistical differences were evaluated by one-way ANOVA followed by Tukey post-hoc test to estimate differences between the separate groups in multiple comparisons. Blue asterisks display the differences compared to the control, while black asterisks represent differences between the different treatment groups. (n.s.: non-significant; *: p<0.05; **: p < 0.01) **(H-I)** Bar graphs show area under the curve (AUC) **(H)** or rate constants (Tau) **(I)** calculations on curves presented in Figure 2G. Scatter plots show the data points from the individual experiments. Data are means ± SEM of five independent experiments, each performed in triplicates. Statistical significance was evaluated by one-way ANOVA followed by Tukey post-hoc test to estimate differences between the separate groups in multiple comparisons. Blue asterisks refer to differences compared to the control, while black asterisks refer to statistical differences between the indicated groups. (n.s.: non-significant; **: p < 0.01). Note that while there is a difference in the AUC values in dynamin 2 knockdowns, due to the smaller PI(4,5)P_2_ decrease in this group, but the τ value (i.e. rate of desensitization) shows no difference. **(J)** Bar graphs show area under the curve (AUC) calculations performed on the data shown in Figure 2H. Cells were treated with the indicated siRNAs and transfected with either empty vector (e.v.) or dominant negative Dynamin2 (K44A). Scatter plots show the results of individual experiments. Data are means ± SEM of seven (control siRNA + e.v. – *blue* and control siRNA + K44A – *red*), five (PIP5K1A siRNA + e.v. – *green* and PIP5K1A siRNA + K44A – *magenta*) or four (EFR3A siRNA + e.v. – *orange* and EFR3A siRNA + K44A – *black*) independent experiments, each performed in triplicates. Statistical significance was evaluated by one-way ANOVA followed by Tukey post-hoc test to estimate differences between the separate groups in multiple comparisons. Blue asterisks refer to the difference compared to the control siRNA + e.v. group, while black asterisks refer to comparisons withing the indicated groups (n.s.: non-significant; ***: p < 0.001, ****: p < 0.0001).

**Figure S2.**
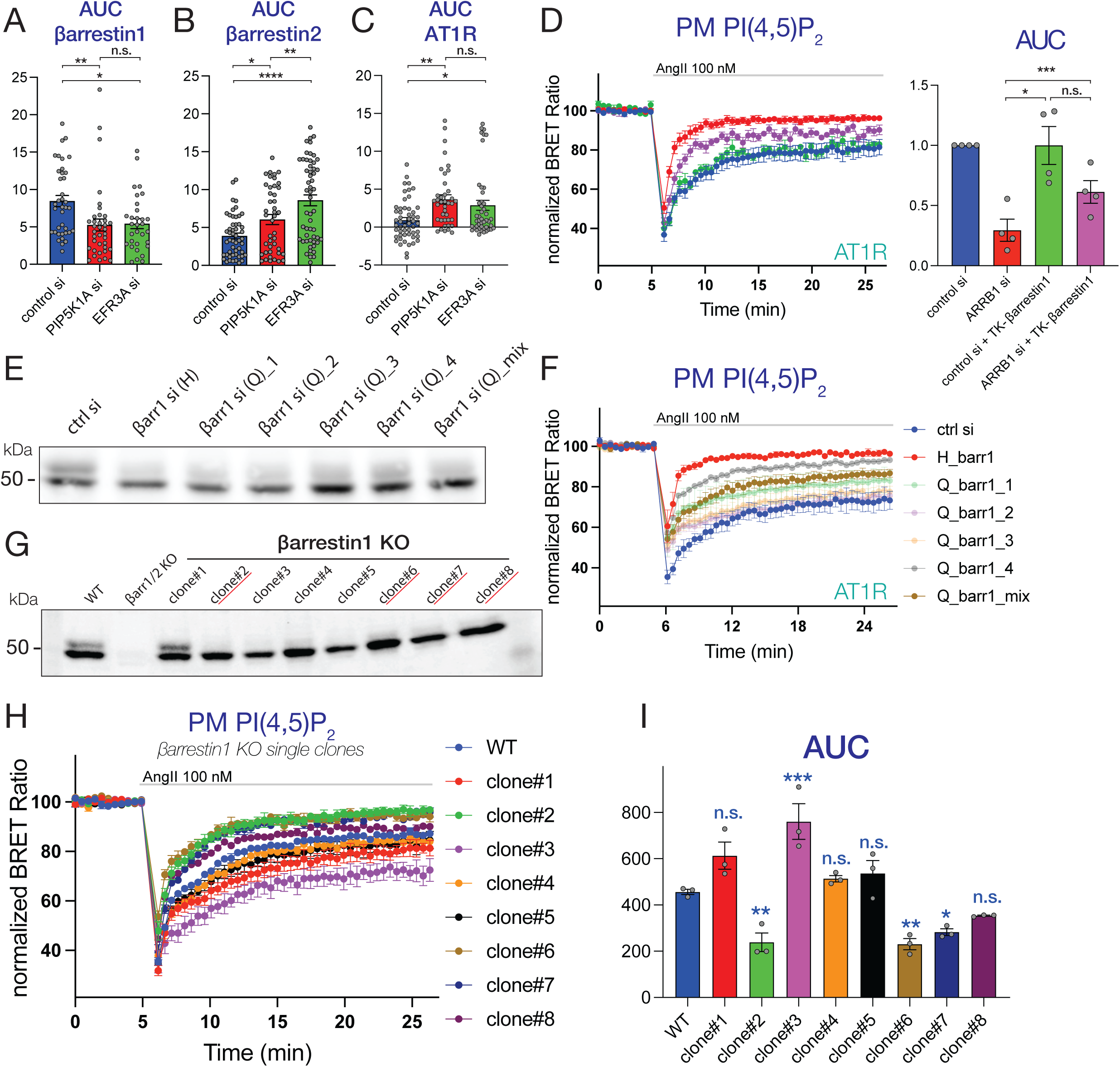
Effects of PIP5K1A or EFR3A knock-down on the PM recruitment of β-arrestins, or clustering of AT1Rs as well as the role of β-arrestin1 on AT1R desensitization. **(A-C)** Bar graphs showing area under the curve (AUC) calculations on data presented in Figure 3 D-F. Cells were treated with the indicated siRNAs and transfected with either TK-promoter driven β-arrestin1-mVenus **(A)**; βarrestin2-mVenus **(B)**; or they stably expressed AT1R-GFP **(C)**. Scatter plots show the results of individual experiments. Data are means ± SEM of 38 (control siRNA – blue), 37 (PIP5K1A siRNA – red) and 31 (EFR3A siRNA – green) cells in panel **A**; 51 (control siRNA – blue), 43 (PIP5K1A siRNA – red) and 54 (EFR3A siRNA – green) cells in panel **B**, obtained in seven independent experiments. For panel **C** the numbers are: 52 (control siRNA – blue), 39 (PIP5K1A siRNA – red) and 46 (EFR3A siRNA – green) from five independent experiments. Statistical significance was evaluated by one-way ANOVA followed by Tukey post-hoc test to estimate differences between the separate groups in multiple comparisons. (n.s.: non-significant; *: p < 0.05, **: p < 0.01; ****: p < 0.0001) (**D**) Rescue experiment performed on HEK-AT1 cells silenced for β-arrestin1 (ARRB1 si) using siRNA-pools and subsequently transfected with TK-promoter driven siRNA-resistant mutants of β-arrestin1. BRET measurements of PM PI(4,5)P_2_ changes after stimulation with 100 nM AngII are shown. (control siRNA – blue, ARRB1 siRNA – red, control siRNA + TK-βarrestin1 – green, ARRB1 siRNA + TK-βarrestin1 – purple). Bar graphs show area under the curve (AUC) calculations to evaluate statistical differences between the groups. Scatter plots show the results of individual experiments. Data are means ± SEM of four independent experiments, each performed in triplicates. Statistical significance was evaluated by two-way ANOVA followed by Tukey post-hoc test to estimate differences between the separate treatment groups in multiple comparisons. Blue asterisks refer to differences compared to control, while black asterisks refer to differences between the indicated groups (n.s.: non-significant; ***: p < 0.001). (E) Representative Western Blots showing the levels of β-arrestin1 and −2 from cell lysates treated with different βarrestin1 specific siRNAs alone or in combination. The upper band is β-arrestin1, the lower one is β-arrestin2. (**F**) BRET measurements of AngII-induced PM PI(4,5)P_2_ changes measured in HEK-AT1 cells treated with different β-arrestin1 targeting siRNAs. After siRNA treatment (4 days in total), cells were transfected with the BRET-based PI(4,5)P_2_ -sensor and stimulated with 100 nM AngII after 5 mins. Data are means ± SEM of three independent experiments, each performed in triplicates. (**G**) Representative Western Blots showing the levels of β-arrestin1 and −2 from cell lysates obtained from wild type (WT), β-arrestin 1/2 double knockout, or β-arrestin1 single knockout clones. The upper band shows β-arrestin1 and the lower band represents β-arrestin2. (**H**) BRET measurements of AngII-induced PM PI(4,5)P_2_ changes measured in HEK-AT1 clones of β-arrestin1 knockouts. Data are means ± SEM of three independent experiments, each performed in triplicates. (**I**) Bar graphs showing area under the curve (AUC) calculations on the curves shown in panel H. To statistically compare the different groups, one-way ANOVA followed by Dunnett’s multiple comparison was used. Scatter plots show the results of individual experiments. Data are means ± SEM of three independent experiments, each performed in triplicates. Significance levels relate to comparison to WT cells (n.s.: non-significant; *: p < 0.05; **: p < 0.01; ***: p < 0.001).

**Figure S3.**
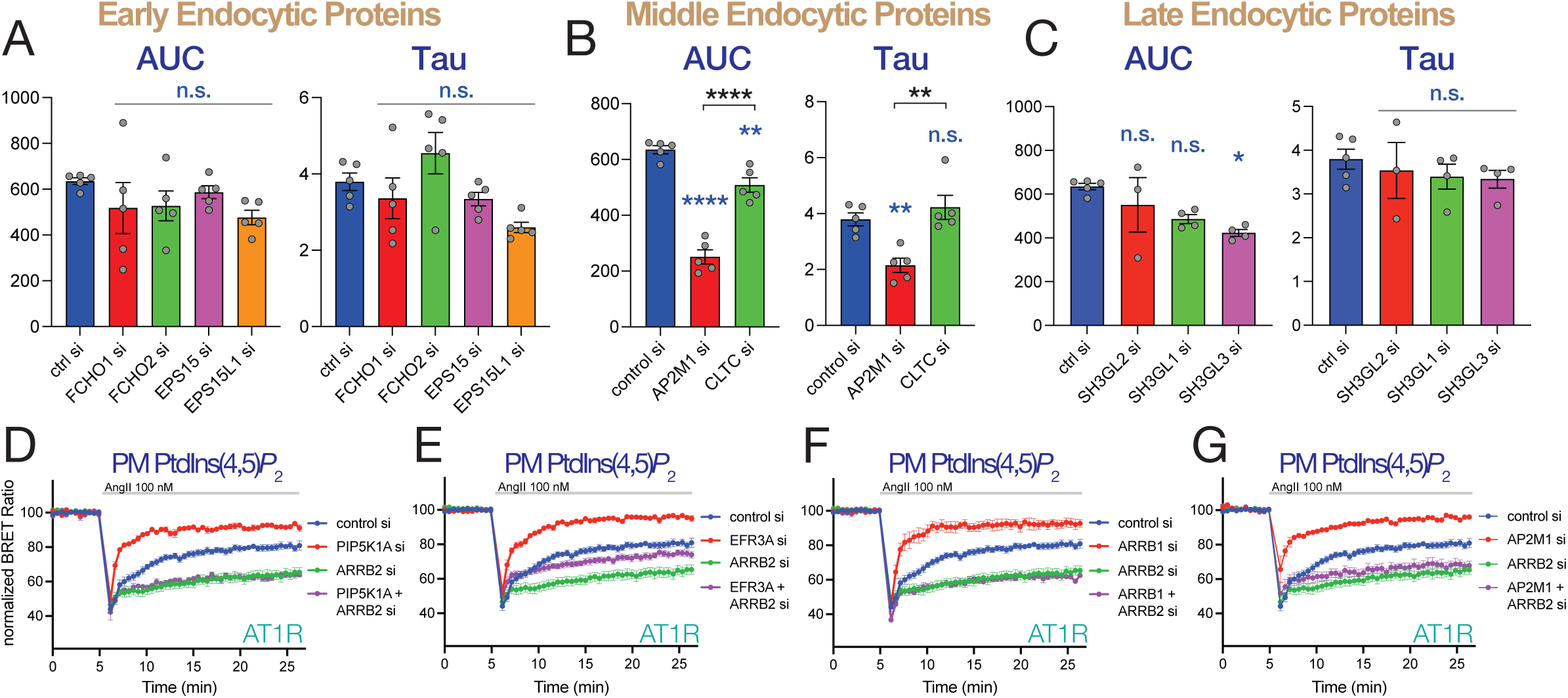
AP2 is required for sustained AT1R activity by supporting βarrestin-1 function. **(A-C)** Bar graphs show area under the curve (AUC) or recovery rate constant (τ) calculations performed on the data shown in Figure 4 B-D. Scatter plots show the results of individual experiments. Data are means ± SEM of five **(A-B)** or four **(C)** (except for the control n=5, and SH3GL2 n=3) independent experiments, each performed in triplicates. Statistical significance was evaluated by one-way ANOVA followed by Tukey post-hoc test to estimate differences between the separate groups in multiple comparisons. Blue asterisks refer to differences compared to control, while black asterisks refer to differences between the indicated individual groups (n.s.: non-significant; **: p < 0.01; ****: p < 0.0001. **(D-G)** BRET analysis of PM PI(4,5)P2 changes in AngII stimulated HEK-AT1 cells after knock-down of PIP5K1A **(D)**, EFR3A **(E)**, β-arrestin1 **(F)**, or AP2M1 **(G)** either alone or in combination with silencing β-arrestin2. Data are means ± SEM of six (control siRNA and ARRB2 siRNA), four (PIP5K1A siRNA, EFR3A siRNA, ARRB1 siRNA, EFR3A + ARRB2 siRNAs, AP2M1 + ARRB2 siRNAs) or three (PIP5K1A + ARRB2 and ARRB1 + ARRB2 siRNAs) independent experiments, each performed in triplicates. *Please note, that for comparison, the same control siRNA treated curves are shown in each panel. Also, ARRB1 siRNA-treated trace in panel **F** is also the same shown in* Figure 3G.

## Additional Supplementary Data

### βarrestin1 siRNA-resistant sequence

(NheI)gctagc*acgcgcacc***atgggcgacaaagggacacgagtgttcaagaaggcaagccccaacgggaaactgacagtgtacc tgggaaagcgggactttgtggaccacattgacctggtggaccccgtggatggcgtggtcctggtggatcctgagtatctcaaagaaaggcgagtctacgtgacactgacctgcgccttccggtatggccgggaagacctggatgtcttgggtctgacttttcgcaaagacctgtttgtggctaacgtgcagtccttcccaccggcccctgaggacaagaagccactgactcggctacaagagcgactcatcaagaagctgggcgagcatgcctaccccttcacctttgagatcccgccaaaccttccgtgctcagtcacattgcaacctgggcctgaggacacagggaaggcctgcggtgtggattatgaagtgaaagccttctgtgctgagaacctggaggagaagatccacaaaaggaattctgtgcggctagtcatccggaaggttcaatatgcccctgagaggcctggccctcagcccacggctgagaccaccagacagttcctcatgtcggacaagcccctgcaccttgaggcatctctagataaagaaatttactatcatggagaacccatcagcgtcaatgtccatgtcaccaacaacaccaacaagactgtgaagaagatcaagatctcggtgcgccagtatgcagacatctgtctcttcaacacagctcagtacaagtgcccagtggccatggaggaagctgatgatactgtggcacccagctcaacattctgcaaggtctacacactgactcccttcctggcaaacaacagagagaagcgggggcttgccctcgacgggaagctcaagcatgaagacacaaatctggcttccagcactctgttgcgggaaggcgccaaccgtgaaatcctgggtatcattgtttcctacaaagtcaaagtgaagctggtggtgtcccggggcggcctgttgggagaccttgcatccagtgacgtggctgtggagcttcctttcactctcatgcaccccaagcctaaagaggagcccccacatcgggaagttccagaatctgaaaccccagtggacaccaatctcatagagcttgacaccaatgatgacgacattgtgtttgaggactttgctcgtcagcggctgaa aggcatgaaggatgacaaggacgaagaggatgatggcaccggctctccacacctcaacaacagatag**gcggccgc(NotI)

---

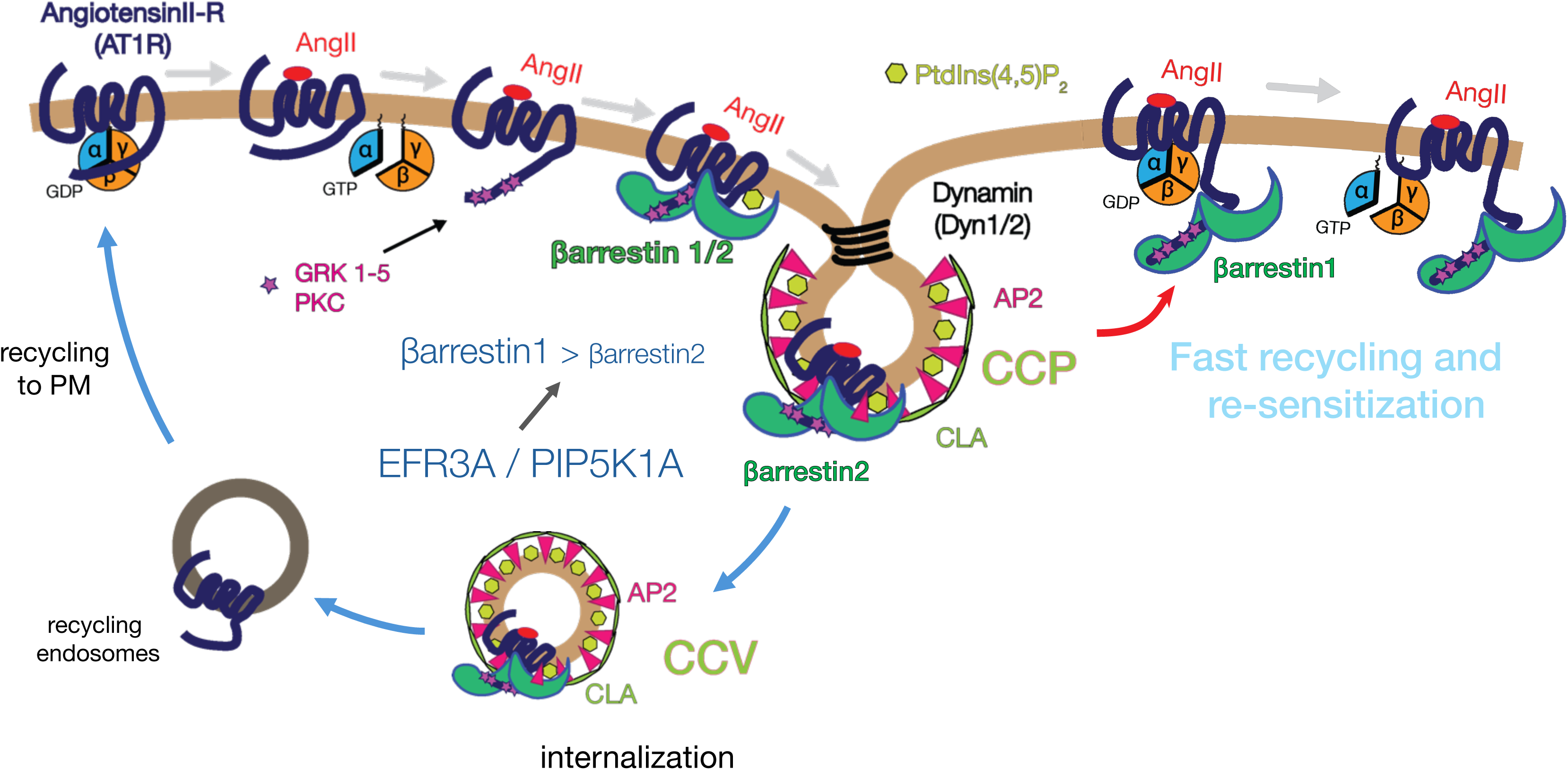

